# A fungal phosphate starvation regulator gates virulence to prioritize nutrient adaptation in response to host phosphate status

**DOI:** 10.64898/2026.03.08.710426

**Authors:** Jacy Newfeld, Seishiro Aoki, Hiromi Haba, Kei Hiruma

## Abstract

Plant-associated microbes must constantly balance environmental adaptation with virulence to survive in fluctuating ecosystems, yet the molecular mechanisms coupling these processes remain poorly understood. Here, we identify *NUTRIENT-DEPENDENT FACILITATOR OF COLONIZATION1* (*NFC1*), a virulence factor in a conditionally pathogenic *Colletotrichum tofieldiae* strain, whose expression is tightly regulated by both temperature and phosphate availability. Leveraging plant phosphate starvation response (PSR) mutants and direct phosphate supplementation to host plants, we demonstrate that *NFC1* expression is driven by the internal phosphate status of the host rather than environmental availability alone. Notably, the fungal PSR regulator *CtPHO4* represses *NFC1* and virulence during phosphate starvation, while simultaneously activating canonical fungal PSR-related genes. Conversely, under phosphate sufficiency, *NFC1* is strongly induced and promotes infection, potentially by modulating the host circadian clock system. This study uncovers a molecular link between metabolic adaptation and virulence programming, providing a conceptual framework for predicting disease dynamics under changing environments.

## INTRODUCTION

In natural and agricultural ecosystems, plants interact with diverse microbes ^1^. These microbes vary in lifestyle, ranging from pathogenic (deleterious) to commensal (neutral) to mutualistic (beneficial). While mutualists provide numerous benefits to their hosts, such as arbuscular mycorrhizal fungi that can provide phosphorus in exchange for photosynthetically derived carbon, pathogens cause severe yield loss annually to crop species around the globe ^2–5^. Lifestyle, however, is not a fixed trait; it shifts over evolutionary time, and it can even vary depending on the host colonized or the environmental conditions ^6–9^. Some pathogens are phylogenetically closely related to mutualists, such as the root-associated species *Colletotrichum tofieldiae* (Ct) ^10,11^. Indeed, despite being sister strains in the Ct phylogeny, strain Ct4 is plant beneficial whereas strain Ct3 is pathogenic on *Arabidopsis thaliana* Col-0 (Arabidopsis) at 22°C under low inorganic phosphate (Pi) conditions. At 26°C, however, Ct3 shifts and becomes plant beneficial in a manner dependent on the Arabidopsis phosphate starvation response (PSR) transcription factors PHOSPHATE STARVATION RESPONSE 1 (AtPHR1) and PHR1-LIKE 1 (AtPHL1) ^10^. A comparable lifestyle switch has been reported for the bacterium *Bacillus amyloliquefaciens*, whose volatile diacetyl causes hyperactivation of the plant PSR under Pi deficiency, but promotes plant growth under Pi sufficient conditions ^12^. The molecular mechanisms driving such lifestyle transitions in response to environmental change remain largely enigmatic.

At the same time as they deal with biotic stresses, like pathogen attack, organisms must cope with abiotic stresses, such as nutrient availability. A major limiting nutrient for plants and fungi alike is phosphorus. The main form of phosphorus taken up by plants and fungi is Pi ^13,14^. To cope with Pi limitation, plants and fungi have, possibly independently, evolved PSRs ^15–18^. In the model plant Arabidopsis, as mentioned above, the PSR is regulated by the transcription factors AtPHR1 and AtPHL1 ^15,19^. Under Pi sufficiency, AtPHR1/AtPHL1 is bound by SPX-domain-containing proteins bound to inositol pyrophosphates (PP-InsPs), and thus, AtPHR1/AtPHL1 cannot bind DNA ^20^. In the model yeast *Saccharomyces cerevisiae*, the fungal PSR is regulated by the transcription factors PHOSPHATE-SENSING TRANSCRIPTION FACTOR 2 (ScPHO2) and PHOSPHATE-SENSING TRANSCRIPTION FACTOR 4 (ScPHO4) ^21^. Under Pi sufficiency, ScPHO4 is phosphorylated, which results in disrupted binding to ScPHO2 and export from the nucleus ^13,22–24^. PHO4 DNA-binding activity is similarly regulated by SPX-domain-containing protein-PP-InsP interactions in at least *Cryptococcus neoformans* and *S. cerevisiae* ^25,26^. It has also been previously characterized that animal pathogenic yeasts rely on the fungal PSR for growth and thus virulence in Pi deficient environments ^27,28^. The virulence of certain plant pathogens has been reported to be phosphate-dependent ^29,30^, suggesting interplay between the plant PSR and plant immunity (as has been proposed previously ^31,32^) and/or between the fungal PSR and virulence. Despite these insights, the role of the PSR in filamentous plant pathogens has yet to be explored. In plant-microbe interactions, microbial effector proteins have been shown to be particularly important for establishment in the host environment, often by modulating plant immunity ^33–35^. Effectors are small, secreted proteins with a short, N-terminal secretion signal known as a signal peptide that targets the protein for the secretory pathway in eukaryotes ^36,37^. While effectors canonically target the host immune system, they have diverse functions in supporting infection, such as facilitating transport of other effectors, targeting the host circadian clock system, or manipulating the host PSR ^38–41^. However, the ways that virulence factors such as effectors respond to the environment and act as intermediaries between environmental stress responses and fungal lifestyle remain unclear.

In this study, by leveraging the drastic lifestyle switch in Ct3 induced by elevated temperature, we first aimed to identify factors that drive this lifestyle switch. We identified a phylogenetically conserved virulence factor, which we here call *NUTRIENT-DEPENDENT FACILITATOR OF COLONIZATION* (*NFC1*). *NFC1*’s expression level, and therefore virulence function, is dependent not only on temperature, but, importantly, also on phosphate availability. We characterize the host response to *NFC1* and determine its possible effect on the host circadian clock. Using a combination of plant and fungal PSR mutants, we find that *NFC1* expression depends both on plant phosphate status, but notably independently of the PSR regulators *AtPHR1*/*AtPHL1*, and on *CtPHO4*, the Ct orthologue of the *S. cerevisiae* PHO regulon regulator *ScPHO4*. Together, these findings provide valuable insights into how plant-associated fungi modulate their virulence in response to fluctuating environmental nutrient conditions.

## RESULTS

### *NFC1* makes a temperature- and phosphate-dependent contribution to virulence

It has been reported that Ct3 can be induced to switch from a pathogenic to a beneficial lifestyle by increasing the temperature from 22°C to 26°C ^10^. To explore the factors driving the lifestyle switch of Ct3 from pathogenic at 22°C to beneficial at 26°C (Fig. 1A-B), we performed *in planta* RNA-sequencing (RNA-seq) at 22°C and 26°C at 12 days post inoculation (dpi) under Pi deficient and sufficient conditions (Fig. 1C). We sampled at 12 dpi because the putative abscisic acid and botrydial biosynthesis cluster (ABA-BOT), previously characterized as contributing to Ct3 virulence, is induced at this timepoint at 22°C under Pi deficient conditions ^10,42^. We first examined the host transcriptome to understand how the plant response to Ct3 can be influenced by the temperature shift. We found that samples clustered more similarly based on temperature than phosphate status, although both factors influenced the transcriptomic response (Fig. S1A). After performing differentially expressed gene (DEG) analysis, we found that there was substantial transcriptional reprogramming dependent on both temperature and phosphate status (Fig. S1B; Table S6-7). We next focused on DEGs at 22°C under Pi deficiency because this is the same condition under which ABA-BOT is induced according to previous reports ^10,42^. We found that Pi starvation and iron homeostasis, factors which themselves are closely linked ^43^, were enriched GO biological process (BP) terms in the set of upregulated DEGs (Fig. S1C). This led us to further investigate phosphate starvation-induced (PSI) gene expression across conditions. We found that PSI gene expression was generally attenuated at 26°C in Ct3 (Fig. S1D). In this way, PSI gene expression seems to negatively correlate with plant growth promotion (PGP) under Pi deficient conditions. This suggests that Ct3 PGP at 26°C is linked to attenuation of the plant PSR, implying an interplay between temperature and phosphate acquisition in plants.

**Figure 1.**
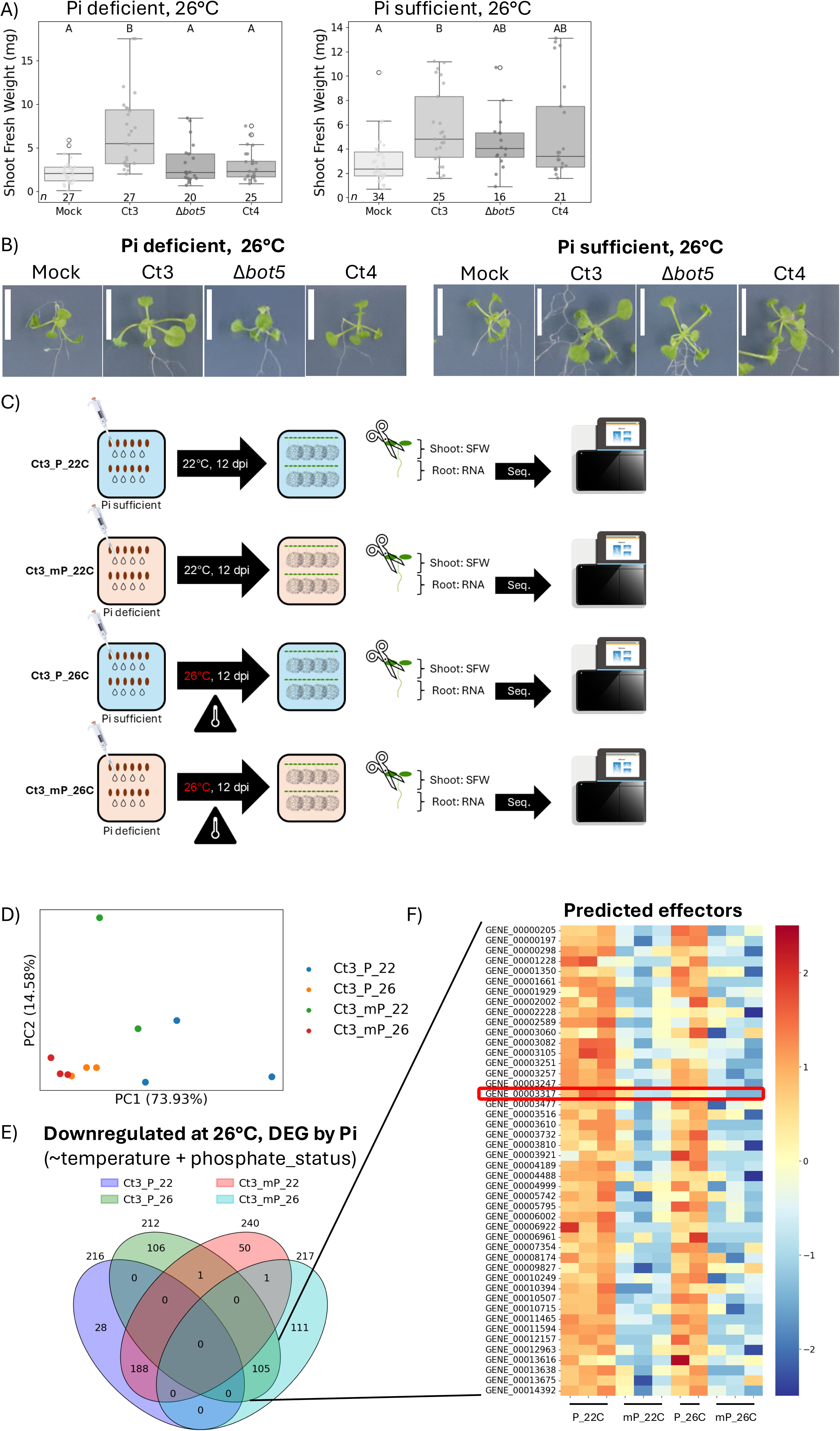
RNA-sequencing reveals *NFC1/EFF3* as a phosphate- and temperature-dependent virulence factor. A) Shoot fresh weight of plants inoculated with fungi at 24 dpi from seed inoculation. Letters above each box represent significance levels from a Dunn test (*p* < 0.05). Numbers below each box represent sample size. B) Representative images of plants quantified in panel A. Bars represent 1 cm. C) Schematic outline of RNA-seq experiment, showing an example of the set-up for Ct3. D) PCA plot of Ct3 transcriptome under sufficient (P) or deficient (mP) phosphate conditions at 22°C and 26°C. E) Four-way Venn diagram of downregulated genes under 26°C compared to 22°C with consideration of phosphate status in the model to identify differentially expressed genes (DEGs). Numbers above each oval indicate the total number of genes in that category. F) Heatmap of all 46 significantly downregulated predicated effectors, with *NFC1* outlined in a red box.

Next, we shifted our focus to the fungal transcriptome. Consistent with a previous report ^10^, ABA-BOT gene expression was reduced in Ct3 at 26°C, whereas interestingly, ABA-BOT gene expression was induced at 26°C in Ct4, especially under Pi deficient conditions (Fig. S2). This is line with abolished Ct4 PGP at 26°C under Pi deficient conditions (Fig. 1A-B). Next, based on our PCA results (Fig. 1D) and the observation that Ct3’s lifestyle switch appears to be dependent on both temperature and phosphate levels, we targeted Ct3 genes that are differentially expressed in response to both factors (Table S4A). This analysis identified 105 genes, notably including numerous candidate effector-coding genes (Fig. 1E, Table S4B). In particular, although predicted effector-coding genes comprise less than 10% of the Ct3 genome, they made up over 40% of the downregulated genes at 26°C compared to 22°C with consideration of Pi status, suggesting an association between the lifestyle switch and attenuation of these candidate virulence factors (Fig. 1E-F). We then generated targeted knockout mutants for a subset of these 46 candidate effector-coding genes, specifically selecting those whose orthologues are not expressed in Ct4 based on a previously published RNA-seq dataset ^10^ (Ct3 locus IDs/NCBI accession = GENE_00000205/KZL71539.1 [*EFF1*, for *EFFECTOR 1*], GENE_00003257/KZL78658.1 [*EFF2*], GENE_00003317/KZL78716.1 [*EFF3*/*NFC1*], GENE_00005795/KZL71948.1 [*EFF4*], GENE_00007354/KZL71142.1 [*EFF5*], GENE_00013616/KZL64564.1 [*EFF6*]). Since these genes are also differentially expressed by environmental phosphate at 22°C, we screened their phenotypes at 22°C under deficient and sufficient Pi (Fig. 2A), including the previously characterized avirulent Δ*bot5* mutant as a control ^10^. This screen successfully identified one key gene, *NFC1*, that consistently contributed to virulence under Pi sufficient, but not Pi deficient, conditions at 22°C (Fig. 2A). This is consistent with its expression profile, where it exhibits an attenuated expression profile under deficient Pi compared to sufficient Pi (Fig. 2B). Furthermore, this coincides with a reduced shoot fresh weight ratio (Mock/Ct3) under Pi deficient conditions at 22°C, indicating a general reduction in Ct3 virulence in this environment (Fig. 2A). Consequently, both wild-type (WT) Ct3 and the Δ*nfc1* mutant induced similar plant growth phenotypes when Pi was deficient. To validate that the Δ*nfc1* mutation is causal to the phenotype we observed at 22°C, we generated complementation lines of *NFC1* in the Δ*nfc1* background driven by its native promoter and terminator (comp-12/22; Fig. S3B). We found that complementation lines expressed *NFC1* to a level comparable to WT (Fig. S3C), and that complementation rescued WT-like virulence at 24 dpi (Fig. 2D). Taken together, these data indicate *NFC1* as a virulence factor whose expression depends not only on temperature but also on phosphate levels.

**Figure 2.**
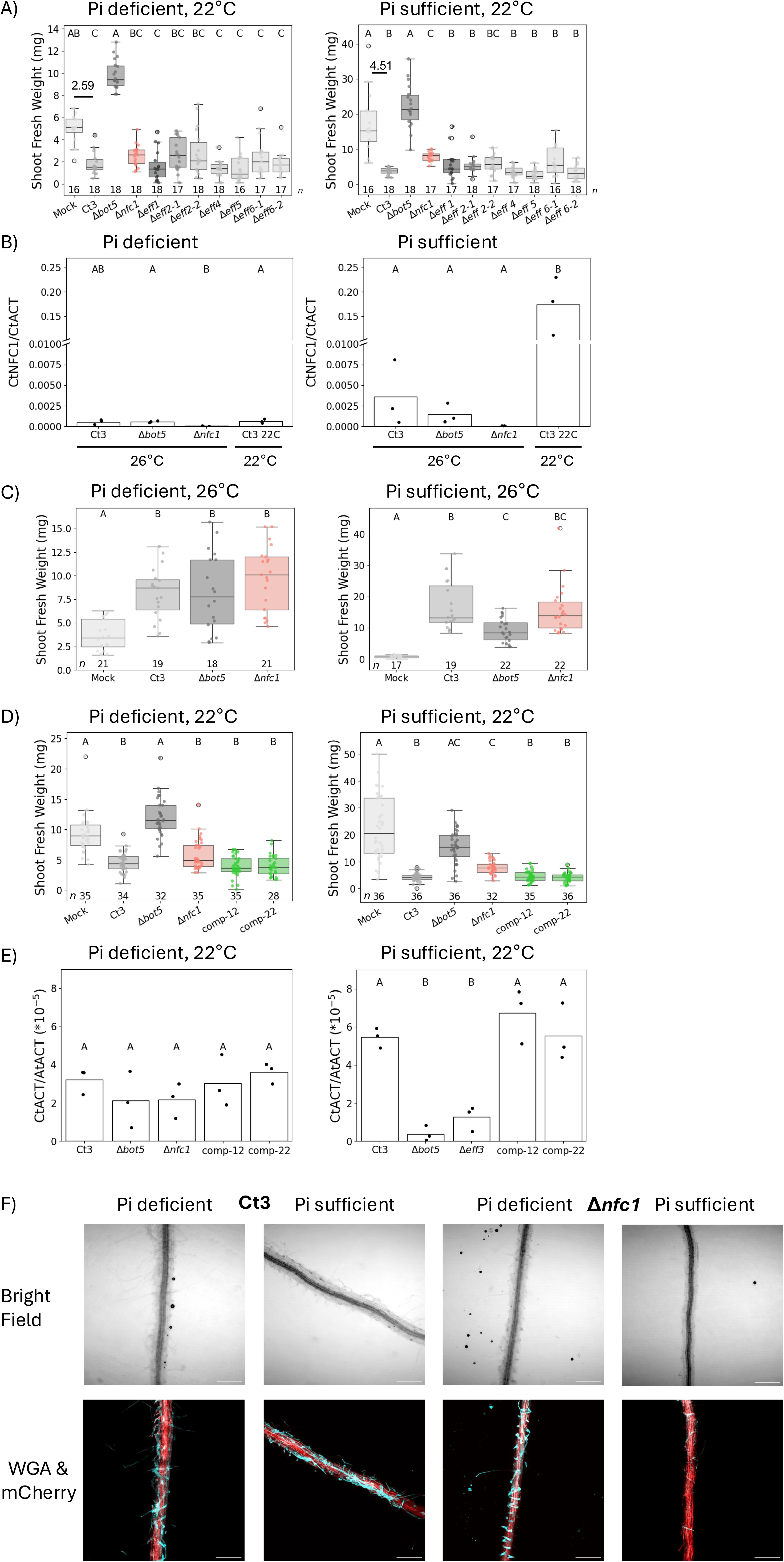
*NFC1* contributes to root colonization under Pi sufficient conditions. A) Shoot fresh weight of plants inoculated with predicted effector knock-out mutants at 24 dpi from seed inoculation. The number above Mock and Ct3 represents the shoot fresh weight ratio (Mock/Ct3). Letters above each box represent significance levels from a Dunn test (*p* < 0.05). Numbers below each box represent sample size. B) Relative expression of *NFC1* under Pi deficient or sufficient conditions at 26°C or 22°C at 2 dpi from direct inoculation. Letters above each bar represent significance levels from a pos-hoc Tukey’s HSD test (*p* < 0.05). C) Shoot fresh weight of plants inoculated with fungi at 26°C at 17 dpi from direct inoculation. Letters above each box represent significance levels from a Dunn test (*p* < 0.05). Numbers below each box represent sample size. D) Shoot fresh weight of plants inoculated with fungi at 22°C at 24 dpi from seed inoculation. Letters above each box represent significance levels from a Dunn test (*p* < 0.05). Numbers below each box represent sample size. E) Root fungal biomass in plants at 24 dpi at 22°C from seed inoculation. Letters above each bar represent significance levels from a post-hoc Tukey’s HSD test (*p* < 0.05). F) Microscopy of WGA-lectin-stained fungal hyphae (cyan) and PIP-mCherry-tagged roots (red) at 4 dpi after direct inoculation with WT Ct3 or Δ*nfc1* under Pi deficient or sufficient conditions at 22°C. Bars represent 200 μm.

### *NFC1* is expressed in plant pathogenic Ct3 at the early infection stage

To better understand the mode of action of *NFC1*, we performed a time-course seed inoculation assay. We started with a seed inoculation assay to get a broad-scale picture of the expression profile under the same conditions used in our preceding experiments and previous studies (see Methods for details). We found that *NFC1* was most strongly induced at 8 dpi, the earliest time point at which we could sample sufficiently colonized roots, under sufficient Pi in Ct3 (Fig. S4A). In contrast, it was not expressed to a high level in plant-beneficial Ct4 or under deficient Pi (Fig. S4A). To ascertain the gene expression profile with higher resolution, we performed a time-course direct inoculation assay, which allows for direct spore attachment to the roots of 7-day-old pre-germinated seedlings. This revealed that *NFC1* was most strongly induced at 2 dpi in Ct3 under Pi sufficient conditions (Fig. S4B). To determine whether the *NFC1* induction observed at 2 dpi is temperature-dependent, we performed a direct inoculation assay with Ct3 and Δ*nfc1* at 26°C. We found that *NFC1* is indeed not strongly expressed at 26°C compared to 22°C and that Δ*nfc1* phenocopies WT Ct3 at 26°C (Fig. 2B-C). These data demonstrate that *NFC1* expression level depends either on the temperature or on the host response to temperature.

### *NFC1* is important for root colonization by Ct3 under Pi sufficient conditions

Interestingly, although Δ*nfc1* had no deficits in growth on rich media or sufficient or deficient Pi media (Fig. S5A-B), the mutant had compromised root fungal biomass at 24 dpi under sufficient Pi conditions, with complementation lines restoring WT-like root fungal biomass (Fig. 2E). Combined with the fact that *NFC1* was induced at an early timepoint, this led us to hypothesize that *NFC1* plays a role in root colonization. To explore this possibility, we analyzed whether Δ*nfc1* mutants had compromised root colonization at the initial colonization process using wheat germ agglutinin (WGA)-lectin-stained fungal hyphae and PIP-mCherry-tagged Arabidopsis. We found that there was a clear reduction in hyphal density on roots inoculated with the Δ*nfc1* mutant compared to WT Ct3 at 4 dpi under sufficient Pi conditions, in line with its contribution to virulence and expression level (Fig. 2F). These data indicate that *NFC1* is indispensable for full root colonization under Pi sufficient conditions. Therefore, we named this gene *NUTRIENT-DEPENDENT FACILITATOR OF COLONIZATION* (*NFC1*).

### *NFC1*’s predicted signal peptide is essential for its virulence function

Secretion of effectors into host cells or the apoplast requires a signal peptide. Using SignalP ^36^, we computationally predicted a putative 24-amino-acid signal peptide at the N terminus of NFC1 (Fig. S3A). To validate its functional consequences, we generated a complementation line in the Δ*nfc1* mutant background with the gene body without its predicted signal peptide (-sec-1/3; Fig. S3B). Although in -sec lines, *NFC1* was expressed to a similar level to WT Ct3 *NFC1*, multiple transgenic lines showed a phenotype similar to the Δ*nfc1* mutant (Fig. S3C-D). This suggests that the predicted signal peptide in *NFC1* is essential for its function in virulence.

### *NFC1* is a phylogenetically conserved, core gene

Next, we sought to understand the evolutionary history of *NFC1*. We therefore constructed a phylogeny based on the sequences of NFC1 using the NCBI whole genome sequence (wgs) and non-redundant protein sequences (nr) databases. This revealed a topology similar to the *Colletotrichum* species tree, indicating a vertical transmission pattern (Fig. 3A; S6) ^44,45^. We then constructed a map of the core and accessory genes based on homologous genes in other Spaethianum species complex members. The results revealed that *NFC1* is a member of the core genome, matching its phylogenetic distribution and high sequence similarity in the Spaethianum species complex (Fig. 3B). These results indicate that *NFC1* is phylogenetically conserved core gene.

**Figure 3.**
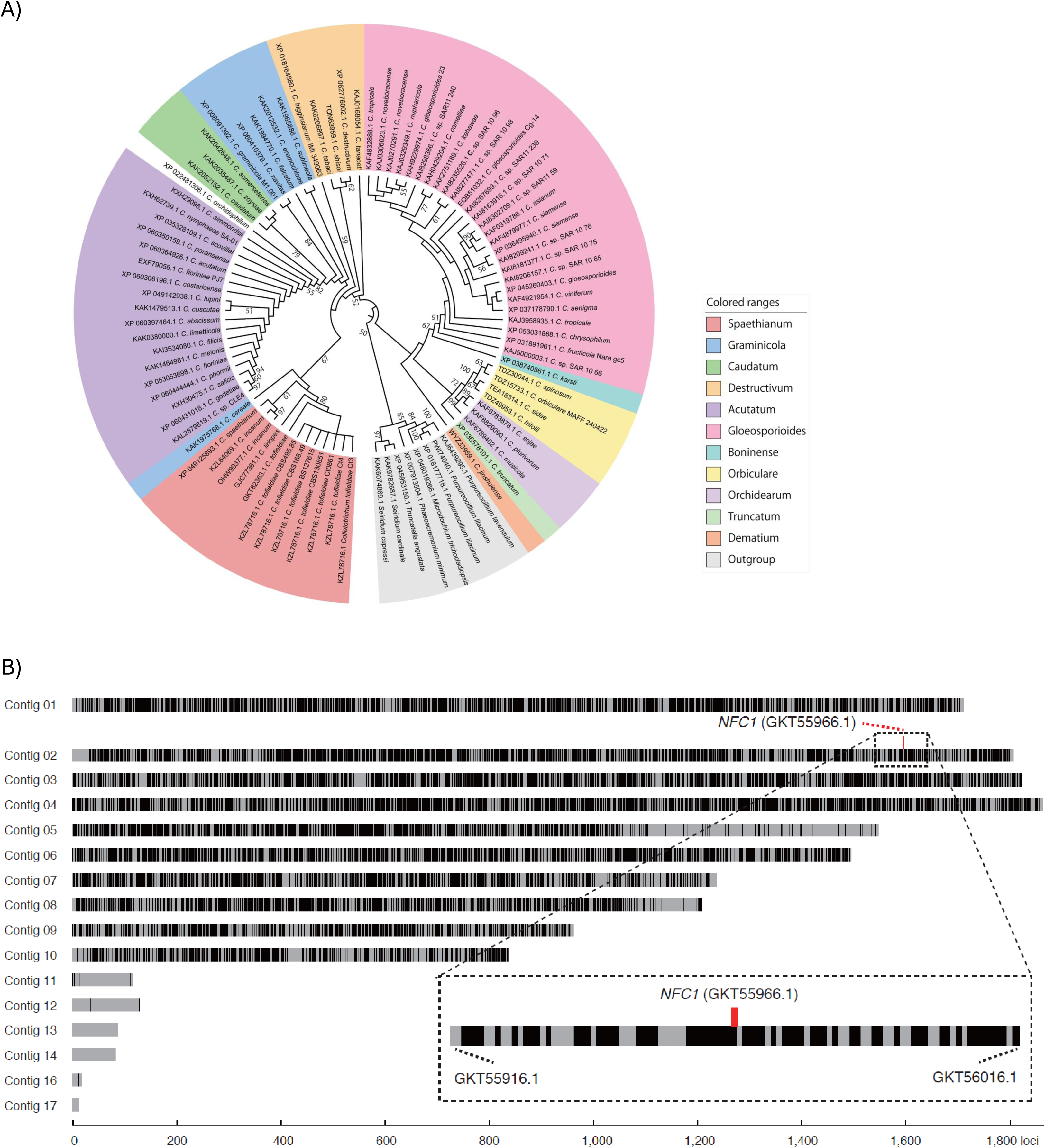
*NFC1* encodes a conserved, core virulence factor. A) Phylogenetic tree showing the relationships among *NFC1* homologs identified from the NCBI whole-genome shotgun (WGS) contigs and the non-redundant (nr) protein database. Colours represent species complexes. The corresponding phylogram including branch length information is shown in Fig. S6. B) Heatmap representing the distribution of core and accessory genes across the 16 contigs of the Ct3 genome. Black bars represent core genes and grey bars represent accessory genes. The red box above contig 02 highlights *NFC1*. Also shown is a close-up of the heatmap, focusing on a region containing 100 genes surrounding *NFC1*.

### *NFC1* may modulate the host circadian clock

To understand the host response to *NFC1* induction at the initial root colonization, we performed RNA-seq at 2 dpi after direct inoculation with Δ*nfc1* or WT Ct3 in Arabidopsis. We found that there was only a marginal shift in the overall Arabidopsis transcriptomic response to WT Ct3 compared to the Δ*nfc1* mutant (Fig. 4A). To investigate the specific host genes that responded to *NFC1*, we examined the DEGs in plants inoculated with Δ*nfc1* compared to WT Ct3 (Fig. 4B-C, Table S8). We found that all DEGs that have been functionally characterized with log_2_FC > 2 and *p_adj_* < 0.05 were related to light and/or circadian rhythms, including *GIGANTEA*, *PRR9*, *ELIP1*, and *PIL6* (Fig. 4B). Because the circadian clock and responses to light were enriched GO terms in the set of Arabidopsis genes induced in response to Δ*nfc1* compared to WT Ct3 (Fig. 4D), we examined the expression of previously characterized circadian clock-related genes (Fig. 4E). We found that there was generally higher expression of circadian clock-related genes in Arabidopsis inoculated with Δ*nfc1* compared to WT Ct3 (Fig. 4E). In line with *NFC1*’s expression profile, some genes showed a similar expression level in WT Ct3-inoculated Arabidopsis under Pi deficient conditions and Δ*nfc1*-inoculated Col-0 under Pi sufficient conditions, including perhaps most notably the circadian clock regulator *CCA1*. Interestingly, *CCA1* has also been proposed as a regulator of the high-affinity phosphate transporter *PHT4;1*, providing a possible mechanistic link between phosphate homeostasis and the circadian clock ^46,47^. Given that PSI genes, including *PHT4;1*, were generally not differentially expressed between the WT and the Δ*nfc1* mutant (Fig. S7), these data suggest that NFC1 may affect the host circadian clock before any potential large-scale gene reprogramming occurs.

**Figure 4.**
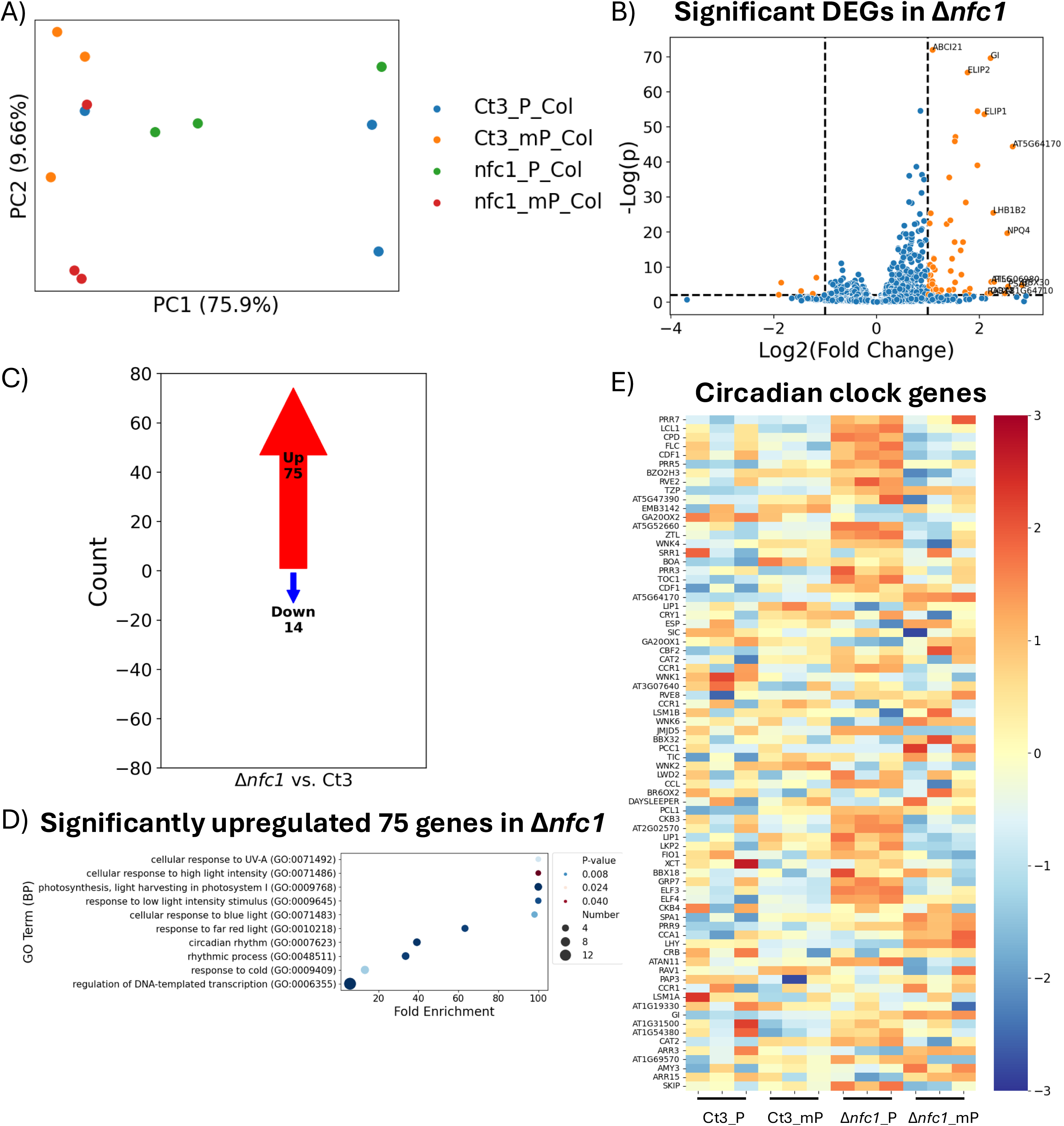
RNA-seq reveals the circadian clock as a potential target of NFC1. A) PCA plot of Arabidopsis transcriptome after inoculation by Ct3 or Δ*nfc1* under Pi sufficient (P) or deficient (mP) conditions at 2 dpi from direct inoculation at 22°C. B) Volcano plot showing significant (*-*Log(*p*) > 2, |Log_2_(Fold Change)| > 1) differentially regulated genes (DEGs; orange) and non-significant genes (blue). Labelled are genes with -Log(*p*) > 2 and |Log_2_(Fold Change)| > 2, or -Log(*p*) > 60. C) Arrow plot showing counts of significant Arabidopsis DEGs (Log(*p*) > 2, |Log_2_(Fold Change)| > 1) after inoculation with Δ*nfc1* compared to WT Ct3. D) Bubble plot of enriched GO terms in the set of 75 significantly upregulated Arabidopsis DEGs in response to Δ*nfc1* inoculation compared to Ct3. E) Heatmap of 78 circadian clock-related genes in Arabidopsis inoculated with Ct3 or Δ*nfc1* under Pi sufficient or deficient conditions, showing induction in Δ*nfc1* inoculation relative to WT Ct3.

### *NFC1*’s contribution to virulence depends on host phosphate status

Because *NFC1*’s expression level and contribution to virulence appear to depend at least on environmental Pi status, we developed three non-exclusive hypotheses about how *NFC1* expression is regulated by the fungus: 1) Ct3 reacts to the environmental phosphate directly; 2) Ct3 reacts to the host PSR; 3) Ct3 reacts to the host internal phosphate status *per se*. To test these hypotheses, we leveraged PSR mutants in Arabidopsis under Pi sufficient and Pi deficient conditions (Fig. S8A-B). In Arabidopsis, the PSR is regulated by the transcription factors PHR1/PHL1 ^15,19^. Pi is taken up into the cell by phosphate transporters, which are trafficked from the endoplasmic reticulum to the plasma membrane by PHOSPHATE TRANSPORTER TRAFFIC FACILITATOR 1 (PHF1)^48^. Once Pi is taken up, it is transported to the xylem by the phosphate transporter PHOSPHATE 1 (PHO1) ^49,50^. Conversely, the ubiquitin-conjugating E2 enzyme PHOSPHATE 2 (PHO2) functions to degrade phosphate transporters, such as the PHOSPHATE TRANSPORTER 1 (PHT1) family and PHO1 ^14,51,52^. At the same time, LOW PHOSPHATE ROOT 1 (LPR1) and LOW PHOSPHATE ROOT 2 (LPR2) function in Pi sensing in the root and suppress primary root growth under low Pi conditions ^53,54^. Therefore, if hypothesis 1 were supported, we would expect to consistently see a contribution to virulence by *NFC1* under Pi sufficient, but not Pi deficient, conditions regardless of plant genotype. If hypothesis 2 were supported, we would expect to see a contribution to virulence even under Pi deficient conditions in PSR-deficient mutant plants such as *phr1phl1* and/or *lpr1lpr2* plants. If hypothesis 3 were supported, we would expect to see an abolished contribution to virulence under Pi sufficient conditions in mutants that experience constitutive internal starvation, such as *phf1*, *pho1*, and/or *pho2*.

We found that the Δ*nfc1* phenotypes on the PSR regulators double mutant *phr1phl1* and the root Pi sensors double mutant *lpr1lpr2* were similar to WT Col-0 under both Pi sufficient and deficient conditions. However, interestingly, Δ*nfc1* had a compromised phenotype in *phf1* plants that are constitutively Pi starved under Pi sufficient conditions (Fig. S8C-E, 5A). To investigate whether the expression level of *NFC1* was also compromised, we performed a direct inoculation on *phf1* plants and found reduced gene expression of *NFC1* in Ct3 inoculated to *phf1* compared to Col-0 plants at 2 dpi under Pi sufficient conditions (Fig. 5B). This suggests that *NFC1* expression is suppressed when plants experience phosphate starvation due to a failure in Pi uptake, rather than through the PSR pathway itself. Next, to address this hypothesis further, we investigated which of root and/or shoot Pi status are important for *NFC1*’s contribution to virulence. Δ*nfc1*’s phenotype on root-Pi-starved *pho2* plants was similarly compromised as *phf1* (Fig. 5C), suggesting that Pi starvation within the specific tissues colonized by Ct3 induces suppression of *NFC1*-mediated virulence. However, surprisingly, the contribution to virulence of *NFC1* was also abolished in shoot-Pi-starved *pho1* plants, although the plants were extremely small even at 17 dpi (i.e. 24-day-old seedlings), meaning that identifying differences between treatments may be difficult for *pho1* plants (Fig. 5C). To test whether localized Pi recovery could modulate *NFC1* expression in the roots, we utilized a cut agar system to apply Pi specifically to either the shoots of *pho1* plants or roots of *pho2* plants (Fig 5D). Based on the data from Thibaud and colleagues ^55^, we applied 100 mM Pi to the shoots of *pho1* plants and roots of *pho2* plants, inoculated fungi to the root tip, then sampled roots at 2 dpi for RNA. We found that exogenous Pi application resulted in significantly increased expression of *NFC1* in both *pho1* and *pho2* plants (Fig. 5E). Similarly, there was a concomitant increase in root fungal biomass (Fig. 5F). Notably, there was no significant increase in *NFC1* expression level nor root fungal biomass under Pi sufficient conditions in WT Col-0 plants (Fig. 5E-F), indicating that there may be an upper limit to Pi content after which *NFC1* expression level remains unchanged. These data demonstrate that *NFC1* expression level depends on plant phosphate status rather than environmental phosphate status (hypothesis 3 above) and imply a shoot-to-root signal induced by shoot Pi starvation.

**Figure 5.**
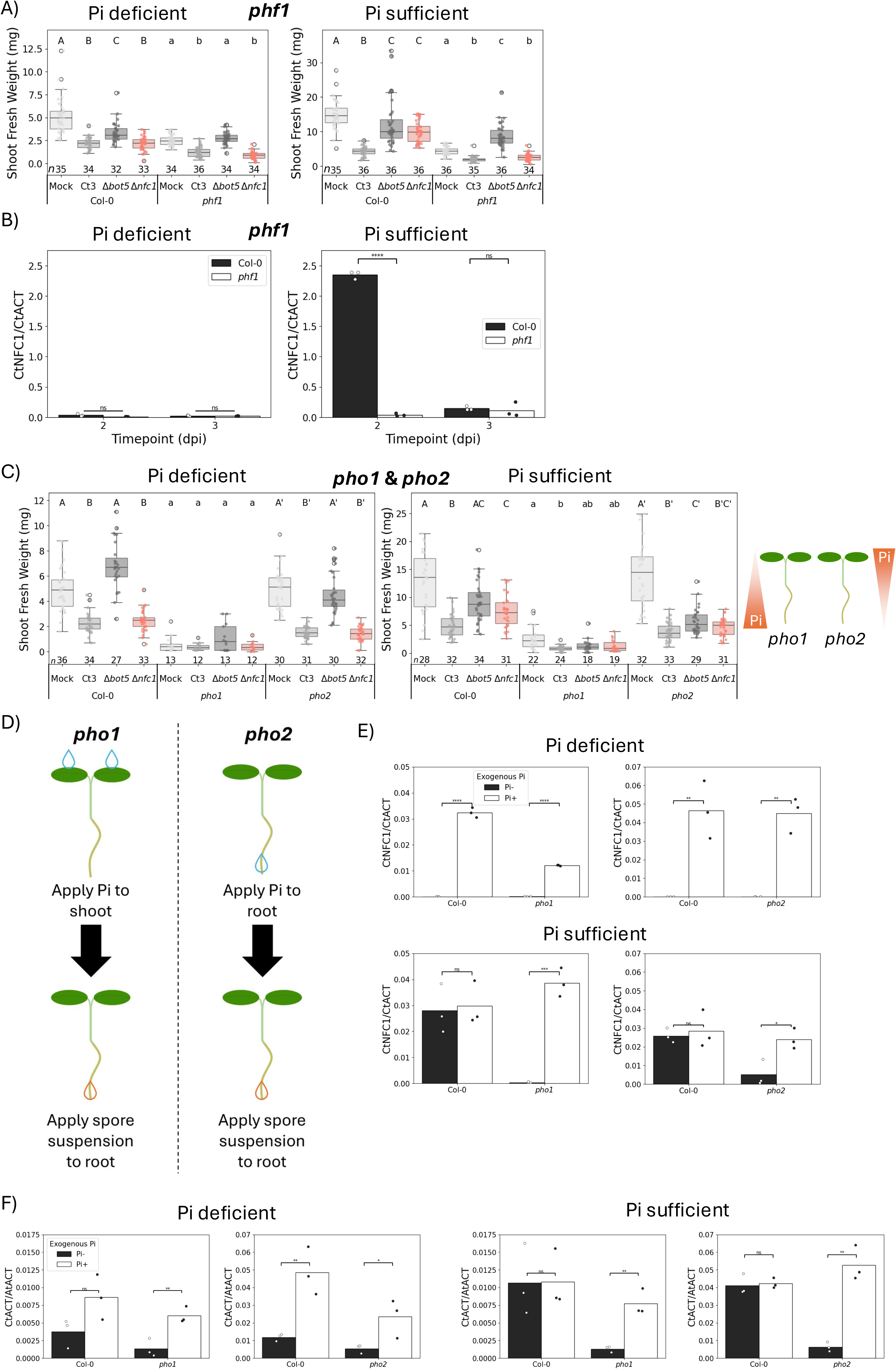
Expression level of *NFC1* is modulated in response to plant phosphate status. A) Shoot fresh weight of Col-0 and *phf1* plants inoculated with fungi at 22°C at 17 dpi from seedling inoculation. Letters above each box represent significance levels from a Dunn test (*p* < 0.05). Numbers below each box represent sample size. B) Relative expression of *NFC1* normalized to *CtACTIN* in Col-0 and *phf1* under Pi deficient or sufficient conditions at 22°C at 2-3 dpi from direct inoculation. Symbols above each bar represent significant differences according to a Student’s independent two-sample t-test (n.s., not significant; ****, *p* < 0.0001). C) Shoot fresh weight of Col-0, *pho1*, and *pho2* plants inoculated with fungi at 22°C at 17 dpi from seedling inoculation. Letters above each box represent significance levels from a Dunn test (*p* < 0.05). Numbers below each box represent sample size. Cartoon beside plots represent Pi content, with *pho1* plants shoot starved and *pho2* plants root starved. D) Schematic outline of exogenous Pi inoculation assay in *pho1* and *pho2* plants. Briefly, 100 mM Pi was applied to shoots of *pho1* or roots of *pho2*, then a spore suspension was applied to the root tip. Roots were sampled at 2 dpi for RNA. E-F) Relative expression of E) *NFC1* normalized to *CtACTIN* or F) *CtACTIN* normalized to *AtACTIN* with and without exogenous Pi application in Col-0, *pho1*, and *pho2* plants at 22°C at 2 dpi after direct inoculation. Symbols above each bar represent significant differences according to a Student’s independent two-sample t-test (n.s., not significant; *, *p* < 0.05; **, *p* < 0.01; ***, *p* < 0.001; ****, *p* < 0.0001).

### *NFC1* expression level and virulence is suppressed by the fungal PSR regulator CtPHO4

Because *NFC1* expression level is responsive to the plant Pi status, we hypothesized that fungal sensing machinery for Pi deficiency may also be important for regulating *NFC1* expression. To explore this possibility, we first identified the orthologous gene to the *S. cerevisiae* PSR regulator *ScPHO4* in Ct3, which we call *CtPHO4* (Ct3 locus ID/NCBI accession = GENE_00002708/GKT55377.1; Fig. S9). In *S. cerevisiae*, ScPHO4 is phosphorylated by the ScPHO85/ScPHO80 cyclin/cyclin-dependent kinase complex and therefore exported from the nucleus under Pi sufficient conditions (Fig. 6A) ^21–24^. Under Pi deficient conditions, ScPHO4 and ScPHO2 together activate the expression of the PHO regulon, which includes, for example, Pi transporters and phosphatases ^21,56^. In this way, while ScPHO2 is a transcriptional co-activator of the PHO regulon, ScPHO4 is the transcription factor that is responsive to Pi status based on its phosphorylation status, which prompted us to focus on *CtPHO4* in subsequent assays.

**Figure 6.**
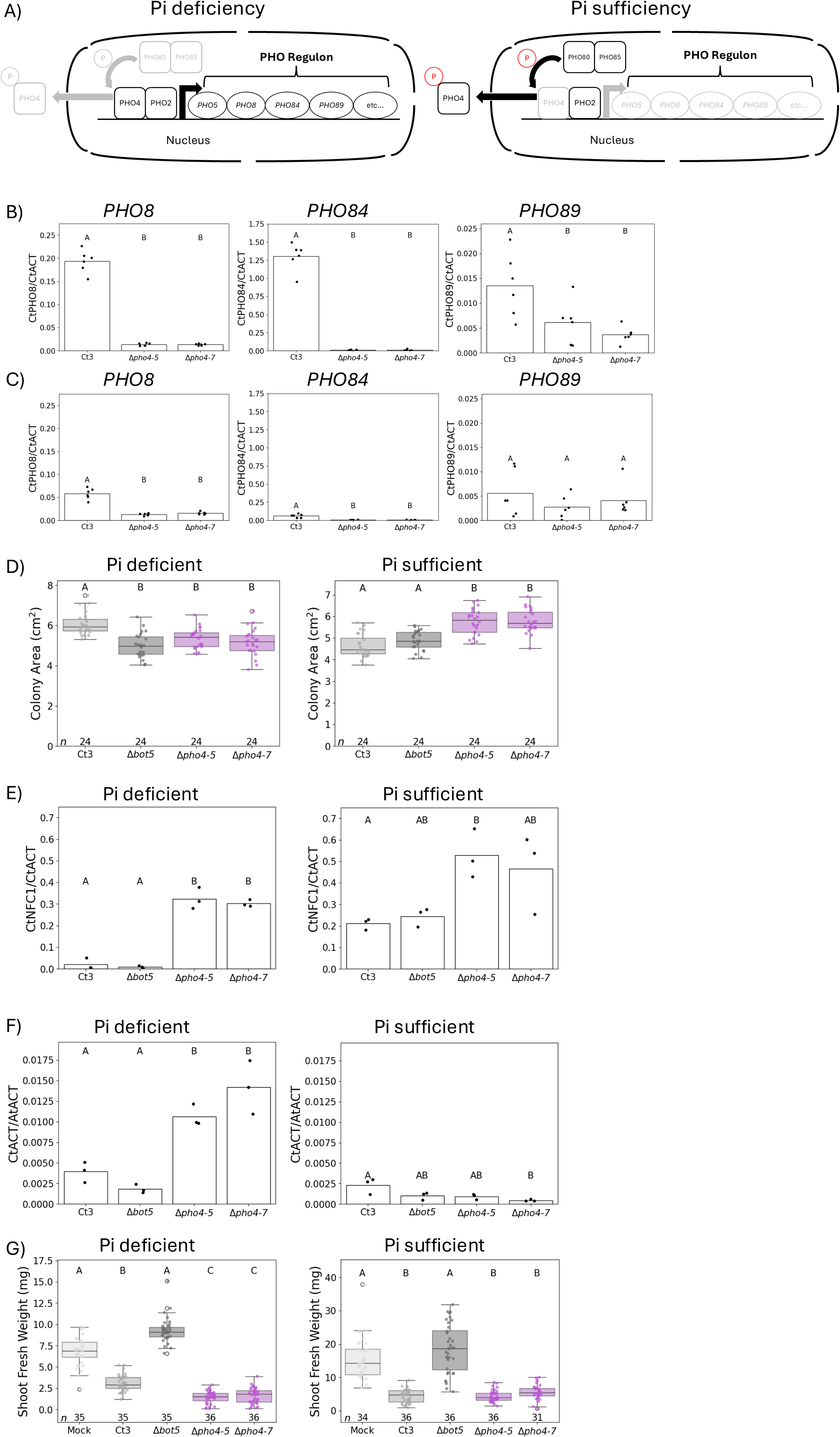
A fungal phosphate starvation regulator modulates expression of *NFC1*. A) Diagram of the PHO regulon. Under Pi sufficient conditions, PHO4 is phosphorylated by the PHO85/PHO80 complex and exported from the nucleus, resulting in non-expression of PHO regulon genes, such as *PHO5*, *PHO8*, *PHO84*, and *PHO89*. Under Pi deficient conditions, PHO4 and PHO2 co-activate PHO regulon genes. B, C) Relative expression of some PHO regulon orthologues in Ct3 normalized to *CtACTIN* in Ct3 and Δ*pho4* fungi from mycelia grown in liquid media under B) Pi deficient or C) Pi sufficient conditions at room temperature for 5 days. Letters above each bar represent significance levels from a post-hoc Tukey’s HSD test (*p* < 0.05). D) Colony area of colonies grown from spores on modified half-strength MS plates with deficient or sufficient Pi at 22°C at 10 dpi. Letters above each box represent significance levels from a Dunn test (*p* < 0.05). Numbers beneath each box represent sample size. E, F) Relative expression of E) *NFC1* normalized to *CtACTIN*, F) *CtACTIN* normalized to *AtACTIN* in Pi deficient and sufficient conditions at 22°C in Ct3, Δ*bot5*, and Δ*pho4* fungi at 2 dpi from direct inoculation. Letters above each bar represent significance levels from a post-hoc Tukey’s HSD test (*p* < 0.05). G) Shoot fresh weight of plants inoculated with fungi under Pi deficient or Pi sufficient conditions at 22°C at 24 dpi from seed inoculation. Letters above each box represent significance levels from a Dunn test (*p* < 0.05). Numbers beneath each box represent sample size.

First, we generated a targeted knock-out of *CtPHO4* in Ct3 to screen its gene expression first to understand its role in the fungal PSR in Ct3 and second to ascertain its role in regulating *NFC1*. To validate that *CtPHO4* is a *bona fide* phosphate starvation regulator in Ct3, we identified orthologues of several PHO regulon genes (Table S1, Fig. 6A). We then grew Ct3 and Δ*pho4* mutant lines in Pi sufficient and deficient liquid media and extracted RNA from mycelia to perform qRT-PCR on several PHO regulon orthologues. This revealed that many of the orthologues were induced by Pi starvation and, crucially, depend on the presence *CtPHO4* (Fig. 6B-C, S10). Importantly, the Δ*pho4* mutant also had compromised growth under Pi deficient conditions (Fig. 6D), but not in rich media (Fig. S5A-B). This suggests first that the PHO regulon genes in Ct3 are indeed Pi responsive, and second, that *CtPHO4* is a regulator of the fungal PSR in Ct3, similar to the role of *ScPHO4* in *S. cerevisiae*. We then monitored the expression level of *NFC1* in Δ*pho4* at 2 dpi and found that multiple Δ*pho4* lines had massively increased expression level of *NFC1*, especially under Pi deficient conditions (Fig. 6E). At the same time, there was increased root fungal biomass under deficient Pi conditions at 2 dpi under Pi deficient conditions, as well as increased virulence at 24 dpi under Pi deficient conditions (Fig. 6F-G). This suggests that the fungal PSR regulated by *CtPHO4* is a critical factor for attenuating *NFC1*-mediated virulence.

## DISCUSSION

In this study, we identified a novel, phylogenetically conserved virulence factor in the conditional plant pathogen Ct3 that contributes to virulence and is expressed in a manner dependent on host phosphate status and temperature, factors which themselves are likely interrelated (Fig. 1, 5). Importantly, the gene’s expression is tightly suppressed by the fungal phosphate starvation regulator *CtPHO4* upon the sensing of Pi starvation by the fungus (Fig. 6). Interestingly, orthologues of *NFC1* were expressed during *C. incanum* infection of Arabidopsis ^57^ and in *C. gloeosporioides* (CgP) infection of Arabidopsis ^58^, potentially indicating a conserved function across *Colletotrichum* pathogens. Moreover, *CgNFC1* was downregulated with co-inoculation of *C. fructicola* (CfE), and there was a concomitant decrease in root fungal biomass ^58^. It is therefore tempting to speculate that *NFC1* may play a similar role in root colonization even in phylogenetically distantly related root pathogens to Ct3 such as CgP.

We found that Δ*nfc1* induces host genes related to the circadian clock system compared to WT Ct3 (Fig. 4). This may indicate that NFC1 affects the host circadian clock. Such manipulation of the circadian clock by virulence factors has been reported, such as the oomycete effector HaRxL10. HaRxL10 was identified in *Hyaloperonospora arabidopsidis* and interacts with CHE to disrupt multiple circadian clock genes as well as salicylic acid-related defence genes to increase disease susceptibility ^40^. There is also an established link between circadian rhythms and immunity ^59,60^. For example, there is periodicity to Arabidopsis resistance to *Pseudomonas syringae*, with this periodicity abolished in plants with mis-expressed *CCA1* or *ELF3* ^61^. Indeed, it was also reported that *CCA1*, in addition to regulating circadian clock genes, also regulates immune-related genes, which the authors hypothesize allows plants to pre-empt their defence response against pathogen attack ^62^. Interestingly, there is also a link between the circadian clock and the plant phosphate transport machinery, with the high-affinity phosphate transporter *PHT4;1* regulated by light and the circadian clock ^46^. Although we did not detect *PHT4;1* as significantly differentially expressed in plants inoculated with Δ*nfc1* compared to WT Ct3 (Table S8, Fig. 4B), a proposed regulator of *PHT4;1*, *CCA1*, was more strongly induced in Δ*nfc1*-inoculated plants than WT Ct3-inoculated plants (Fig. 4E). This possibly supports the idea of an interrelated network of immunity, the circadian clock, and phosphate acquisition that together are involved in the host response to *NFC1*.

While conditional pathogenicity has previously been attributed to evolutionary genetic factors, such as loss of accessory chromosomes, there has been relatively less attention given to how fungi adjust their lifestyles in response to the environment ^63–65^. Nevertheless, it has been established that temperature can impact microbial lifestyle, with previous literature having largely focused on the phenomenon of increased virulence in pathogens in response to elevated temperature ^66–69^. In line with this, we observed that Ct4, which is beneficial at 22°C under Pi deficient conditions, behaves plant neutrally at 26°C, and at the same time, induces expression of ABA-BOT genes (Fig. 1A-B, S2B). Conversely, Ct3, which is pathogenic at 22°C, promotes plant growth at 26°C, and at the same time, reduces expression of ABA-BOT genes (Fig. 1A-B, 2C, S2A). This system therefore presents a model with which to understand conditional, temperature-dependent pathogenicity. Using this system, we identified a novel virulence factor whose expression level is modulated not only by the temperature, but also by the plant phosphate status and fungal phosphate starvation machinery (Fig. 5-6). Although the detailed mechanism remains to be elucidated, there is possibly crosstalk between the plant PSR and temperature, as we have shown in Ct3-inoculated plants (Fig. S1D) and as has been evidenced by a previous report in Ct3 ^10^. In that paper, Ct3 promoted plant growth at 26°C in WT Col-0 plants, but notably not *phr1phl1* plants, suggesting that the host PSR is an important determinant of fungal lifestyle under mildly elevated temperature conditions. Similarly, a previous report identified that Pi content in Arabidopsis was decreased at 28°C compared to 22°C, suggesting an interaction between temperature and Pi acquisition ^70^. In line with this, we have shown that both host phosphate status and temperature influence the expression of a fungal virulence factor (Fig. 1, 2B, 5). Collectively, these data underscore how plants and their associated fungi fine-tune their genetic programs in response to the environment.

Previous studies have also demonstrated the role of phosphate in the plant-pathogen interaction. In response to excess Pi, when challenged by *Magnaporthe oryzae*, rice (*Oryza sativa*) experienced increased susceptibility ^29^. Similarly, a more recent study demonstrated that Pi accumulation leads to suppression of hydrogen peroxide accumulation in rice leaves, resulting in increased susceptibility to *M. oryzae* ^75^. In line with these findings, it was also found that Pi deficiency increases resistance of cotton (*Gossypium hirsutum*) to *Verticillium dahliae* ^30^. Similarly, Ct3 has reduced pathogenicity under Pi deficient conditions, which coincides with a reprogramming of virulence, as evidenced by the Pi-dependency to *NFC1* expression level (Fig. 2B, S4, 5B, 5E). In support of this, it was previously demonstrated that Ct3 induces chlorosis under Pi sufficient, but not Pi deficient, conditions ^10^. This implies that the fungus senses Pi deficiency, and that such sensing is important for virulence programming.

Modulation of virulence by single virulence factors has been reported numerous times in plant-pathogen interactions, but it has been historically less studied how fungi reprogram specific virulence factors in response to environmental conditions ^40,71–73^. The roles of the virulence factor we identified seems to be distinct from previously characterized NUDIX effectors, which induce the host phosphate starvation response by hydrolyzing PP-InsPs that regulate host Pi homeostasis ^41^. Instead, *NFC1* is responsive to host phosphate status, but this response does not appear to depend on the PSR regulators *PHR1*/*PHL1* nor the root phosphate sensors *LPR1*/*LPR2* under Pi sufficiency (Fig. 5, S8D-E). This is expected, given that these factors function primarily during Pi deficiency. However, in PSR mutants that exhibit Pi-starved phenotypes either in shoots or roots even under Pi sufficiency, *NFC1* is suppressed, and exogenous Pi application to the Pi deficient tissues is sufficient to rescue the expression level comparable to the WT plants (Fig. 5E). It is therefore possible either that the balance of shoot and root Pi is important or that local Pi deficiency in either tissue is sufficient to modulate *NFC1* expression level. The latter appears more likely, as Pi application to Col-0 plants under Pi sufficient conditions did not significantly alter *NFC1* expression level. The precise shoot-to-root molecular signal remains to be uncovered, but could possibly depend on small RNAs, such as microRNA399, which moves from shoot to root in Arabidopsis in response to Pi deficiency ^74^. This represents a previously uncharacterized phenomenon in which filamentous fungi sense host phosphate status, even in uncolonized tissues, as a pathway to reprogramming virulence.

The PHO regulon has been well-studied using model yeasts, but the role of the PHO regulon in filamentous fungi has remained elusive ^16,27,76^. Here, we demonstrate that *CtPHO4*, the orthologue of the yeast PHO regulon regulator *ScPHO4*, suppresses pathogen virulence at least partly through attenuation of the expression level of *NFC1* (Fig 6E-G). This is distinct from the role of the PHO regulon in animal pathogenic yeasts, where its positive contribution to virulence is considered a consequence of adaptation to Pi-limiting conditions ^27,28^. In Ct3, *CtPHO4* suppresses the full virulence of Ct3, likely functioning to balance virulence gene expression with phosphate utilization gene expression. Therefore, there may be some functional diversification of the PHO regulon in plant-associated filamentous fungi such as Ct3 compared to yeasts. In filamentous fungi, it is possible this strategy of balancing virulence with the fungal PSR is necessary to prioritize PSR gene expression under Pi deficiency in order to maximize the intracellular Pi content and adaptation to Pi deficient environments, a mechanism that would parallel the role of *AtPHR1* in balancing immunity and growth in Arabidopsis ^31,32,77^. In this hypothesis, *CtPHO4* would be necessary for growth under Pi limitation. In support of this, Δ*pho4* mutants had decreased growth under Pi deficient conditions (Fig. 6D), suggesting that activation of the PHO regulon is necessary for optimal growth under Pi deficient conditions. Interestingly, in *S. cerevisiae*, the PHO regulon is responsive to the plant hormone strigolactone, with strigolactone inducing the fungal PSR not only in *S. cerevisiae*, but also in filamentous fungi such as *Fusarium graminearum* ^78^. This may represent a strategy in which plants target the fungal PSR to reduce the virulence of fungal pathogens. Our report on *CtPHO4* represents the first study, to the best of our knowledge, to provide genetic evidence that filamentous fungal nutrient starvation responses shape the virulence program.

In summary, we have identified *NFC1* as a novel virulence factor whose regulation is governed by the interplay between fungal phosphate sensing and plant phosphate status. The tight coordination of fungal virulence with nutrient adaptation mechanisms highlights the remarkable plasticity of plant-associated microbes, which can shift between pathogenic and beneficial states depending on host and/or environmental factors ^10,42^. Strikingly, in the beneficial strain Ct4, *NFC1* is induced by elevated temperature (Fig. S2B), in direct contrast to Ct3, where its expression is suppressed. Unraveling the functional link between abiotic stress adaptation and virulence regulation will facilitate the development of sustainable agricultural strategies to mitigate pathogenicity while harnessing beneficial plant-microbe interactions.

## METHODS

### Plant and Fungal Growth Conditions

All plant materials were in the *A. thaliana* Col-0 background. Arabidopsis genotypes included Columbia 0 (Col-0; WT), *phf1* ^48^, *phr1phl1* ^15^, *lpr1lpr2* ^79^, *pho1* ^50^, and *pho2* ^52^. Fungal materials were in the Ct3 (*Colletotrichum tofieldiae* MAFF 712333) background ^10^. Ct4 (*Colletotrichum tofieldiae* MAFF 712334) was also used in the 26°C assay ^10^. Fungal cultures were maintained on Mathur’s media plates with 3% (w/v) agar for short-term storage (2 weeks or less), or in 25% glycerol, 75% water-spore suspension (v/v) for long-term storage (more than 2 weeks).

### Fungal Inoculations

#### Seed Inoculation

Seed inoculation was performed as described previously described ^10,42^. Briefly, Arabidopsis Col-0 seeds were sterilized two times with 70% ethanol for 30 seconds, then 6% sodium hypochlorite with 0.01% Triton X for 5 minutes. Seeds were then washed 5 times with sterile distilled water (DW). Sterilized seeds were sown onto modified half-strength Murashige and Skoog (MS) square plates with 1.3% (w/v) INA Agar BA-10 (INA Food Industry) (pH = 5.1) containing either 625 μM KH_2_PO_4_ (Pi sufficient condition) or 50 μM KH_2_PO_4_ (Pi deficient condition) (see ^10^ for details). Fungal conidia were washed from a Mathur’s media plate using 5 mL sterile DW, rinsed once with sterile DW, then counted in a hemocytometre. Conidial suspensions were diluted to 5,000 spores/mL, then 3 μL (∼15 spores) was inoculated 3 cm below seeds (Fig. 1C). Plates were sealed with surgical tape and incubated vertically in a growth chamber at 22°C or 26°C, ∼40% humidity, and a 10-hour photoperiod (∼100 μmol m^−2^ s^−1^). At 24 dpi, shoots were manually separated from roots and weighed for shoot fresh weight measurements. At the indicated timepoint, roots were collected, flash frozen in liquid nitrogen, and stored at -80°C until DNA or RNA was extracted.

#### Seedling Inoculation

Because some Arabidopsis genotypes have a low germination rate, we used 7-day-old pre-germinated seedlings for assays involving Arabidopsis mutants. Arabidopsis seeds were sterilized as outlined above and inoculated to modified half-strength MS media containing either sufficient or deficient Pi. Seeds were incubated in a growth chamber at 22°C or 26°C, ∼40% humidity, and a 10-hour photoperiod for 7 days. 7-day-old Arabidopsis seedlings were then transplanted from half-strength MS plates to new modified half-strength MS plates of the same Pi condition using disinfected forceps. Conidia were harvested, washed, and diluted as above, then inoculated 2 cm beneath the root tip. At 17 dpi, shoots were manually separated from roots and weighed for shoot fresh weight measurements. At the indicated timepoint, roots were collected, flash frozen in liquid nitrogen, and stored at -80°C until DNA or RNA was extracted.

#### Direct Inoculation

To facilitate assays on the early infection stage between Ct and Arabidopsis, we used a direct inoculation method to allow for direct spore attachment to the roots of 7-day-old pre-germinated seedlings. 7-day-old Arabidopsis seedlings were transplanted from half-strength MS plates to new half-strength MS plates of the same Pi condition. Conidia were harvested, washed, and diluted as above, then inoculated directly onto the root tip. At the indicated timepoint, shoots were manually separated from roots and weighed for shoot fresh weight measurements and roots were collected, flash frozen in liquid nitrogen, and stored at -80°C until DNA or RNA was extracted.

#### Exogenous Pi application

Inoculation was performed as for Direct Inoculation (above) with modifications (Fig. 5D). First, the agar was cut 1 cm from the top edge of the plate such that, when sown, seedling shoots were aerial while the roots were on agar. This functioned to physically separate the shoot from the root such that application of phosphate was specific to either shoot or root. After transplanting, 100 mM KH_2_PO_4_ (Pi+) or 100 mM KCl (Pi-) was applied to the shoots (2 μL to each cotyledon, *pho2* plants) or roots (5 μL to root tip, *pho1* plants) of 7-day-old seedlings. Conidia were prepared and inoculated as above. At 2 dpi, roots were flash-frozen in liquid nitrogen and stored at -80°C until RNA was extracted.

### Cloning & Transformation

#### Cloning

For knock-out generation, primers (Table S2) were designed to amplify 1500 bp regions immediately upstream and downstream of the gene of interest. Regions were amplified from the wild-type Ct3 genome by PCR using PrimeSTAR HS DNA Polymerase (TaKaRa), following manufacturer instructions. To isolate the region, PCR products were electrophoresed on a 1% agarose gel at 100 V for ∼30 minutes, then fragments of the appropriate size were cut from the gel. DNA was isolated using the FastGene Gel/PCR Extraction Kit (NIPPON Genetics) following manufacturer instructions. Vectors were constructed with upstream and downstream regions using *Escherichia coli* Stellar Competent Cells (Clontech), the In-Fusion HD Cloning Kit (TaKaRa), and the pBIG4MRHrev plasmid containing hygromycin (hyg) and kanamycin antimicrobial resistance genes, following manufacturer instructions. Successful transformants were verified by colony PCR. Plasmids were isolated from successful transformants using the NucleoSpin Plasmid EasyPure miniprep kit (Machery-Nagel). For knock-in, sequenced and successfully constructed plasmids were transformed to *E. coli* Stellar Competent Cells, then pDNA was isolated using the NucleoBond Xtra Midi kit (Machery-Nagel).

For complementation, primers were designed to amplify the gene of interest, its upstream promoter (∼2000 bps) and its downstream terminator (∼300 bps). Regions were amplified by PCR, then isolated as above. Vectors were constructed with the region as above, except using pPZPnat1, which contains kanamycin and nourseothricin (NRS) resistance genes, instead of pBIG4MRHrev. Successful transformants, plasmid isolation, and verification proceeded as above, using NRS instead of hyg.

#### Transformation: Knock-Out

For targeted gene knock-out for fungi, the protocol was performed as described previously ^10^. Briefly, *Agrobacterium tumefaciens* str. C58C1 was transformed with the construct by electroporation at 1800V. Transformed *Agrobacterium* cells were cultured in liquid LB supplemented with kanamycin overnight, then cells were washed with GI broth containing 200 μM acetosyringone twice. Fungi were cultured in flasks with 100 mL Mathur’s agar for one week prior to the transformation, then conidia were washed off with GI broth. Conidia were subsequently washed twice with GI broth containing 200 μM acetosyringone. *Agrobacterium* cells (OD_600_=0.4) and fungal conidia (10^7^ spores/mL) were mixed in a 1:1 ratio, then plated on a sterile paper filter affixed to a GI agar plate containing 200 μM acetosyringone. Hygromycin-resistant colonies were selected by transferring filter paper to Mathur’s plates supplemented with hyg, cefotaxime (cefo), and spectinomycin (spec), then removed from plates and discarded. Plates were incubated until fungal colonies emerged, which were then transferred to a fresh Mathur’s plate containing hyg, cefo, and spec. Fungal colonies were subsequently re-streaked onto fresh Mathur’s with hyg, cefo, and spec, then genotyped by colony PCR to validate the gene replacement.

#### Transformation: Knock-In

For gene knock-in (complementation) of fungi, the protoplast-polyethylene glycol (PEG) protocol was performed as described previously with minor modifications ^42^. Briefly, spores of Ct3 were harvested from 250 mL Mathur’s liquid media and inoculated to fresh 250 mL Mathur’s liquid media for 48 hr at 60 rpm for germination. Germinated spores were collected, and cell walls were digested using driselase (Sigma-Aldrich), yatalase-plus (TaKaRa), and beta-glucanase (Sigma-Aldrich). Protoplasts were purified, washed, then 10-15 μg of pDNA was added alongside 40% PEG. Transformants were selected on potato dextrose agar (PDA; Difco) with 0.6 M glucose and 200 μg/mL nourseothricin (NRS). Colonies were re-streaked to Mathur’s with 200 μg/mL NRS, then genotyped by colony PCR to validate insert presence.

### Fungal Growth Assay

#### Without Plants

Agar plugs (ϕ = 5 mm) of 7-day-old fungal cultures on Mathur’s media plates were transferred to new Mathur’s media plates. Colony size was examined after 3 days of growth by measuring edge to edge divided by two in ImageJ/Fiji ^80^.

#### With Plants

Inoculation was performed as described above (Seed Inoculation). Fungal colony size was determined 10 dpi by Fiji by measuring the colony area.

### DNA Extraction

To analyze root fungal biomass, DNA was extracted from flash-frozen roots inoculated with fungi. First, roots were homogenized by a Shake Master NEO (Biomedical Science). DNA was then extracted using the cetyltrimethylammonium bromide (CTAB) method as previously described ^81^. DNA was subsequently stored at - 20°C.

### RNA Extraction & Gene Expression Analysis

#### RNA Extraction

To analyze gene expression, flash-frozen roots or mycelia stored at -80°C were homogenized by a Shake Master NEO (Biomedical Science). Total RNA was extracted using the NucleoSpin RNA kit (Macherey-Nagel), according to manufacturer instructions. From this RNA, cDNA was synthesized from 100-200 ng total RNA using the PrimeScript RT Reagent Kit (TaKaRa).

#### Quantitative Reverse Transcription PCR (qRT-PCR)

cDNA was amplified in THUNDERBIRD Next SYBR qPCR Mix (TOYOBO) with 250 nM primers using a CFX Opus 384 Real-Time PCR System (BIORAD) in a volume of 10 μL. The average of two technical replicates is shown per biological replicate.

#### RNA Sequencing & Analysis

500-1000 ng of total RNA per sample was sent to Rhelixa (Japan) for sequencing, including 3 biological replicates per sample. After Poly-A selection by NEBNext Poly(A) mRNA Magnetic Isolation Module, the strand-specific libraries were generated by NEBNext Ultra II Directional RNA Library Prep Kit. These libraries were sequenced by an Illumina NovaSeq X Plus platform (150 bp x 2 paired-end reads), resulting in approximately 40 M pairs per sample for the 26°C vs. 22°C experiment or 20 M pairs per sample for the Δ*nfc1* vs. Ct3 experiment.

The obtained reads were first quality-checked by FastQC ^82^. From the temperature assay, sample Ct3_mP_22_3 was found to have virtually no fungal reads, so it was excluded from subsequent analysis. Reads were then aligned to the Ct3, Ct4, or Arabidopsis genomes (TAIR10) using hisat2 (v2.1.0) ^83^. Generated SAM files were sorted into BAM files using samtools (v1.10) ^84^. featureCounts (v2.0.0) was then used to obtain gene counts ^85^. Genes with a lower count than 5 across all treatments were excluded from further analysis. Based on these counts, DEGs were analyzed in Python (v3.8.10) using PyDESeq2 ^86^. Using PyDESeq2, we modeled both phosphate status and temperature in a multi-factor analysis to call DEGs based on phosphate status and temperature, allowing statistical testing with consideration of both variables. DEGs were considered significant if |log_2_FC| >1 and *p*_adjusted_ < 0.05. Predictions of secreted proteins in the Ct3 predicted proteome were performed using SignalP 6.0 ^36^, with candidate effectors then predicted with EffectorP 3.0 ^37^. Proteins were considered predicted to be secreted when the probability was greater than 0.9, and predicted secreted proteins were considered to be predicted effectors when the probability was greater than 0.9 and there was no predicted transmembrane helix based on TMHMM ^87^. Heatmaps were drawn using the heatmap function in seaborn based on normalized gene counts ^88^. For plant genes, related genes were pulled from the Arabidopsis Information Resource (TAIR) using the Gene search function, followed by manual inspection of gene descriptions ^89,90^. Specifically, for phosphate-related genes, we performed a search using the TAIR Gene search function for gene names containing the term “phosphate”, yielding 121 genes. For circadian clock-related genes, we used the TAIR Gene search tool for gene key words containing the term “circadian clock”, yielding 78 genes. For plant gene GO analysis, the TAIR GO Term Enrichment tool was used. For comparison with the previously reported RNA-seq of Ct3 and Ct4 ^10^, genes were considered not expressed in Ct4 if fragments per kilobase per million reads (FPKM) values were less than 10.

### Microscopy

Inoculations were performed as described above (Direct Inoculation) using Δ*nfc1* or wild-type Ct3 fungi and PIP-mCherry tagged Arabidopsis. At 4 dpi, roots were harvested and immediately fixed in 4% paraformaldehyde phosphate buffer solution (PFA; FUJIFILM) for 1 hr at room temperature. PFA was replaced with ClearSee solution, and roots were incubated overnight at 4°C. ClearSee was replaced with wheat germ agglutinin (WGA) conjugated to an FITC solution (Sigma-Aldrich) at a concentration of 5 μg/mL in ClearSee. Roots were incubated with WGA stain overnight at 4°C, then washed twice with ClearSee, mounted on a slide, and visualized by FITC signal (487-545 nm) for fungal hyphae or mCherry signal (570-670 nm) for plants on a confocal microscope (OLYMPUS FLUOVIEW FV3000) using a 10x UPLXAPO objective lens.

### *NFC1* Phylogenetic Tree

Sequences similar to *NFC1* were collected using a BLASTP search against the GenBank NR and whole-genome shotgun contigs databases the NCBI whole-genome shotgun (WGS) contigs and the non-redundant (nr) protein database in February 2026. Highly divergent (E-value > 1.00E-10 or query cover < 50%) sequences were excluded from the dataset. Sequence alignment was performed with MAFFT (v7.2211) using the - -auto option ^91^. Phylogenetic analysis was conducted using IQ-TREE (v2.4.0; options: - m MFP -T AUTO -b 100) ^92^.

### Ct3 Core Genome Analysis

Genome analysis was performed using the OrthoMarkov clustering method implemented in the GET_HOMOLOGUES package (v14112024) with default parameters ^93^. Homologous genes from 7 genomes of the *C. spaethianum* species complex (Table S3) were identified and clustered using the script compare_clusters.pl, which uses BLASTP for pairwise gene comparisons. The resulting clusters were organized into a genome matrix to represent gene presence and absence patterns. Subsequent analysis of the genome matrix was performed using the parse_pangenome_matrix.pl utility script included in the GET_HOMOLOGUES package. Genes were categorized into four groups based on their distribution across genomes: core (present in all genomes), soft core (present in most genomes), shell, and cloud genes. Shell and cloud genes represent intermediate- and low-frequency accessory genes, respectively ^94^. To visualize the distribution of core (core and soft core) and accessory (shell and cloud) genes, a heatmap was generated for the 16 contigs of the C3 genome containing genes that are translated into amino acids (Figure S3). This plot highlights the spatial organization of genes, with distinct patterns for core and accessory components in the genome.

### PHO4 Sequence Alignment

*CtPHO4* was identified by BLASTp analysis using ScPHO4 (NCBI accession: DAA12477.1) as the query and the Ct3 predicted proteome as the subject. CtPHO4 structure was predicted by AlphaFold3 ^95^. The alignment between CtPHO4 and ScPHO4 was performed using Clustal W ^96^, then alignment was visualized by FoldScript ^97^.

### Statistical Analysis

All statistical analysis and data visualization was performed in Python (v3.8.10) using the following libraries: pandas ^98,99^, seaborn ^88^, statsmodels ^100^, SciPy ^101^, and scikit-posthocs ^102^. Data with 3-6 data points (i.e. qPCR data) were visualized as bar plots, and data with more than 6 data points (i.e. colony size, shoot fresh weight data) were visualized as box plots with outliers represented as closed circles. Assumptions for parametric tests were validated using the Shapiro-Wilk test for normality and Levene test for homoscedasticity. Differences among means or medians were then analyzed using either one-way ANOVA or Kruskal-Wallis test, respectively. Differences between samples were then analyzed by post-hoc Tukey’s Honest Significant Differences (HSD) or a Dunn test, respectively. To correct for multiple comparisons, the Bonferroni method was applied. For qRT-PCR and qPCR, relative gene expression and root fungal biomass were analyzed by the double delta Ct method.

## ACKNOWLEDGEMENTS

We wish to thank members of the Hiruma Lab—namely Momoko Takagi, Yuniar Devi Utami, Tan Anh Nhi Nguyen, Ren Ujimatsu, Akito Shiina, Xuan Xuan, and Risa Ayano—for fruitful discussion and commentary on this work. We thank Javier Paz-Ares for *phr1phl1*, *phf1*, and *pho2* seeds, Yves Poirier for *pho1* seeds, and Thierry Desnos for *lpr1lpr2* seeds. This work was supported by a JSPS grant (grant number JP23K26903 to KH), JST SPRING grant (grant number JPMJSP2108 to JN), JST ALCA-Next grant (grant number JPMJAN23D4 to KH), and a JST FOREST grant (grant number JPMJFR200A to KH).

## DATA AVAILABILITY

Raw RNA sequencing reads have been deposited in the DDBJ (PRJDB40195 for the temperature-related RNA-seq [Fig. 1], PRJDB40210 for the host response to *NFC1* [Fig. 4]). The authors do not report original code.

## AUTHOR CONTRIBUTIONS

KH initiated the project. JN and KH directed the research. JN and HH conducted the experiments. JN conducted the RNA-seq analysis. SA performed the core and accessory genome analysis and constructed the phylogenetic tree. JN analyzed the data. JN and KH wrote the manuscript, with feedback from all other authors.

## SUPPLEMENTARY TABLES

**Table S1.** PHO regulon orthologues in Ct3 as identified by BLASTP.

| Gene symbol | NCBI reference protein sequence | Ct3 locus ID | E-value |
| --- | --- | --- | --- |
| *PHO4* | NP_116692.1 | GENE_00002708 | 3e-08 |
| *PHO5* | NP_009651.3 | GENE_00012685<sup>1</sup> | 2e-39 |
| *PHO12* | NP_012087.1 | GENE_00012685<sup>1</sup> | 1e-39 |
| *PHO11* | NP_009434.1 | GENE_00012685<sup>1</sup> | 1e-39 |
| *VTC1* | NP_010995.1 | GENE_00007825 | 1e-54 |
| *VTC2* | NP_116651.1 | GENE_00004693<sup>2</sup> | 3e-143 |
| *VTC3* | NP_015306.1 | GENE_00004693<sup>2</sup> | 1e-154 |
| *VTC4* | NP_012522.2 | GENE_00002463 | 5e-142 |
| *PHO86* | NP_012418.1 | GENE_00001514<sup>3</sup> | 0.59 |
| *PHO89* | NP_009855.1 | GENE_00013372 | 8e-153 |
| *PHO8* | NP_010769.3 | GENE_00001477 | 6e-169 |
| *PHO84* | NP_013583.1 | GENE_00003000 | 0.0 |

<sup>1</sup> These genes were merged into *CtPHO5*.
<sup>2</sup> These genes were merged into *CtVTC2*.
<sup>3</sup> This gene was excluded from further analyses due to low similarity.

**Table S2.**
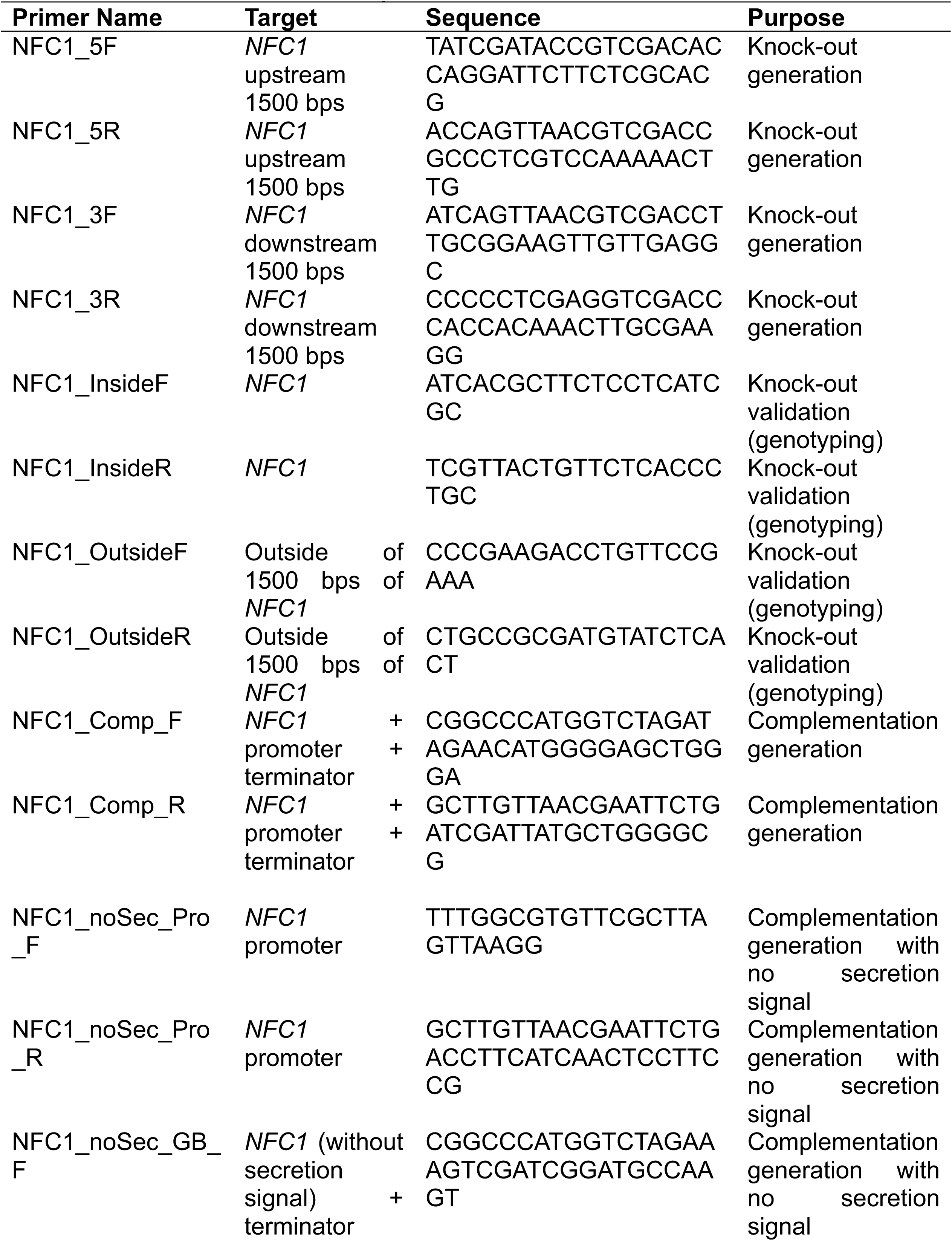
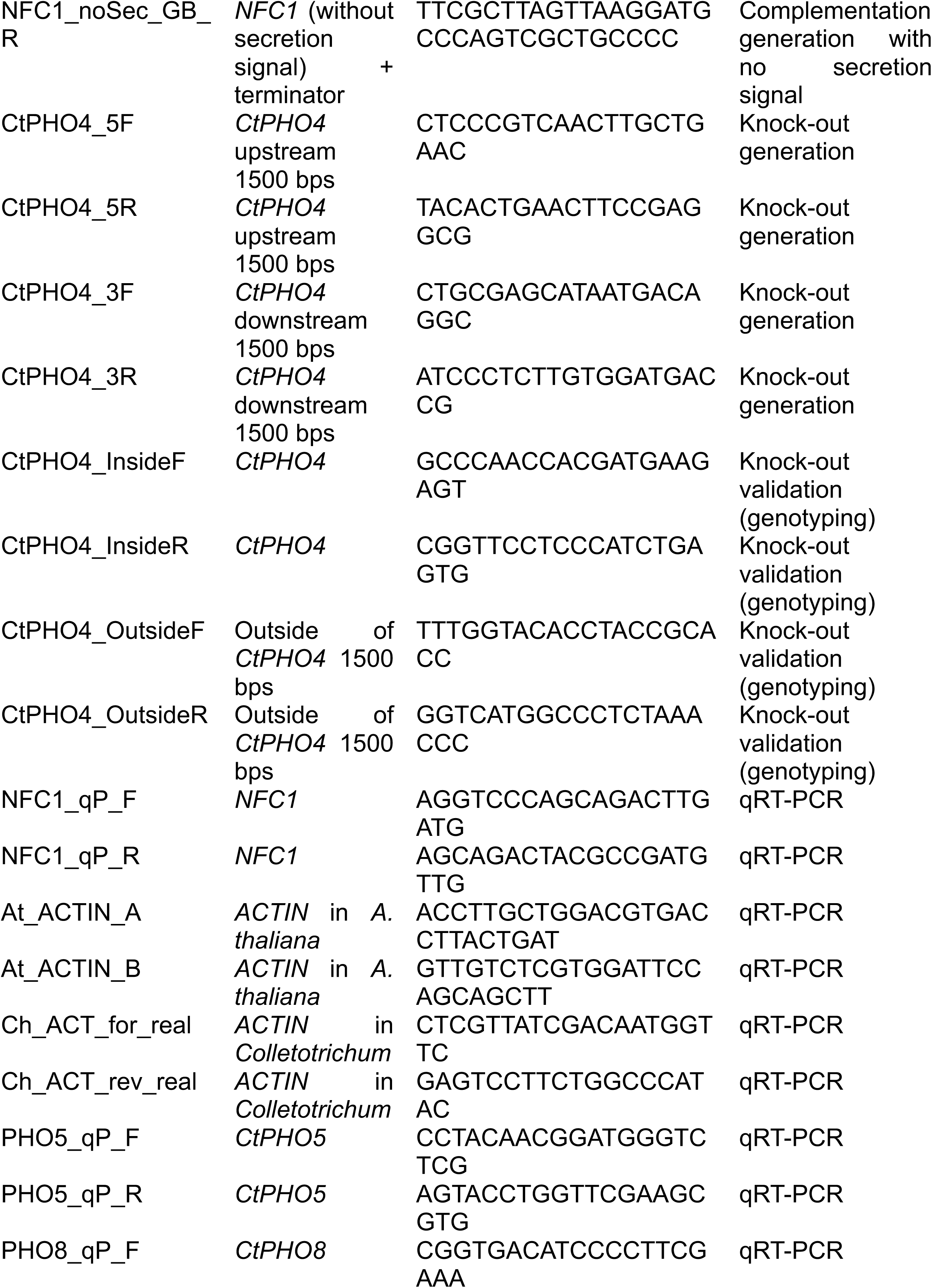
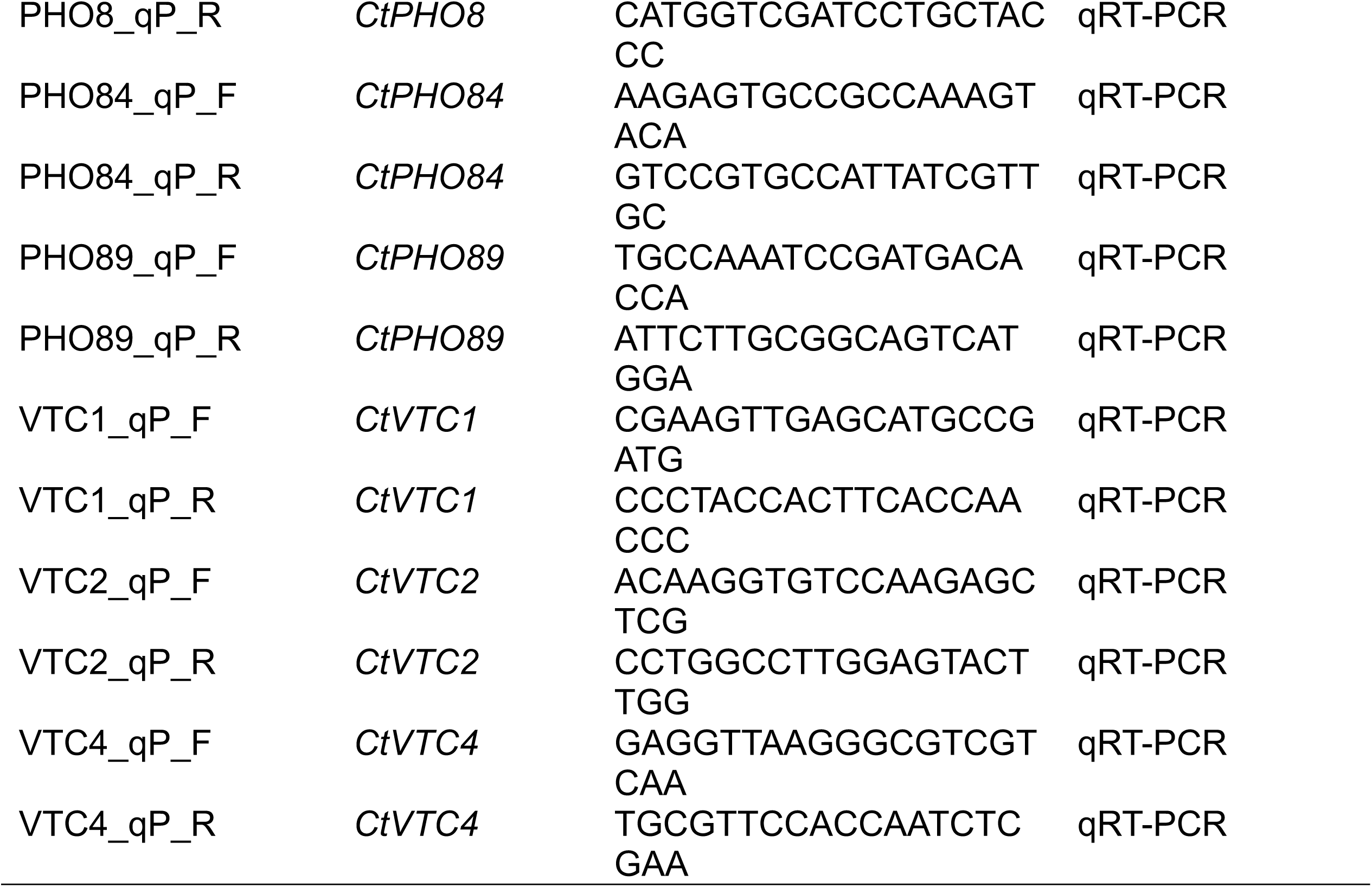
Primers used in this study.

**Table S3.** List of strains used in the genome analysis for prediction of core and accessory regions in Ct3.

| Strains | Assembly Accession |
| --- | --- |
| *Colletotrichum tofieldiae* MAFF 712333 (Ct3) | GCA_022836555.1 |
| *Colletotrichum tofieldiae* MAFF 712334 (Ct4) | GCA_022836575.1 |
| *Colletotrichum tofieldiae* 0861 (Ct61) | GCA_022836595.1 |
| *Colletotrichum incanum* MAFF 238704 | GCA_001625285.1 |
| *Colletotrichum incanum* MAFF 238712 | GCA_001855235.1 |
| *Colletotrichum liriopes* MAFF 242679 | GCA_022179045.1 |
| *Colletotrichum spaethianum* MAFF 239500 | GCF_022836535.1 |

## SUPPLEMENTARY FIGURE LEGENDS

**Figure S1.**
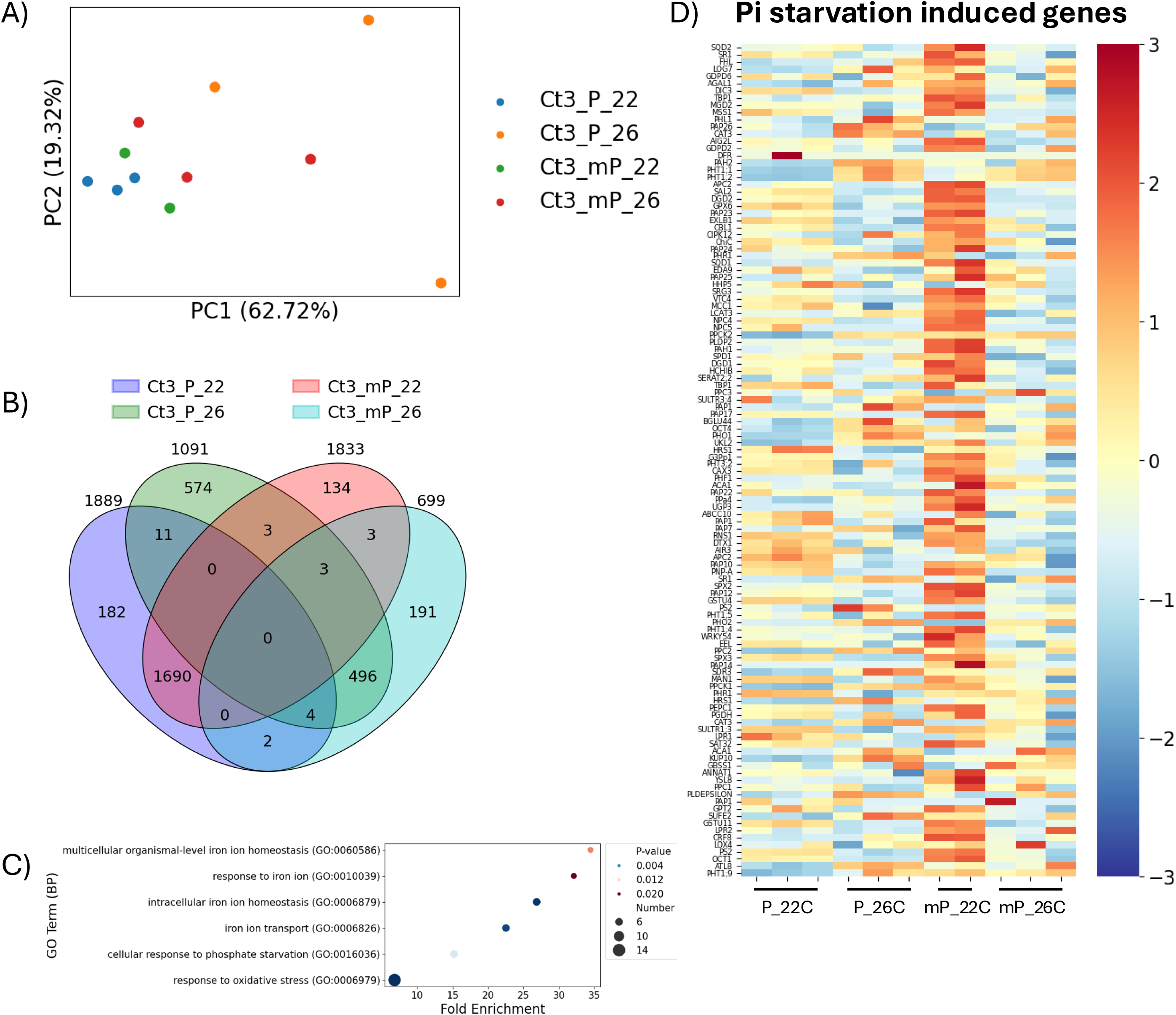
Plant growth promotion negatively correlates with PSI gene expression. A) PCA plot of Arabidopsis transcriptome at 12 dpi after seed inoculation with Ct3 at 22°C or 26°C under Pi sufficient (P) or deficient (mP) conditions. B) Venn diagram showing significantly upregulated Arabidopsis DEGs (Log(*p*) > 2, Log_2_(Fold Change) > 1) in response to inoculation by Ct3 at 22°C or 26°C under Pi sufficient (P) or deficient (mP) conditions. Numbers above each oval represent the total number of significantly upregulated DEGs in that condition. C) GO enrichment analysis of 134 upregulated Arabidopsis DEGs in response to inoculation with Ct3 at 22°C under Pi deficient conditions. D) Heatmap of Arabidopsis phosphate starvation-induced genes at 26°C or 22°C under Pi sufficient or deficient conditions in Ct3-inoculated plants.

**Figure S2.**
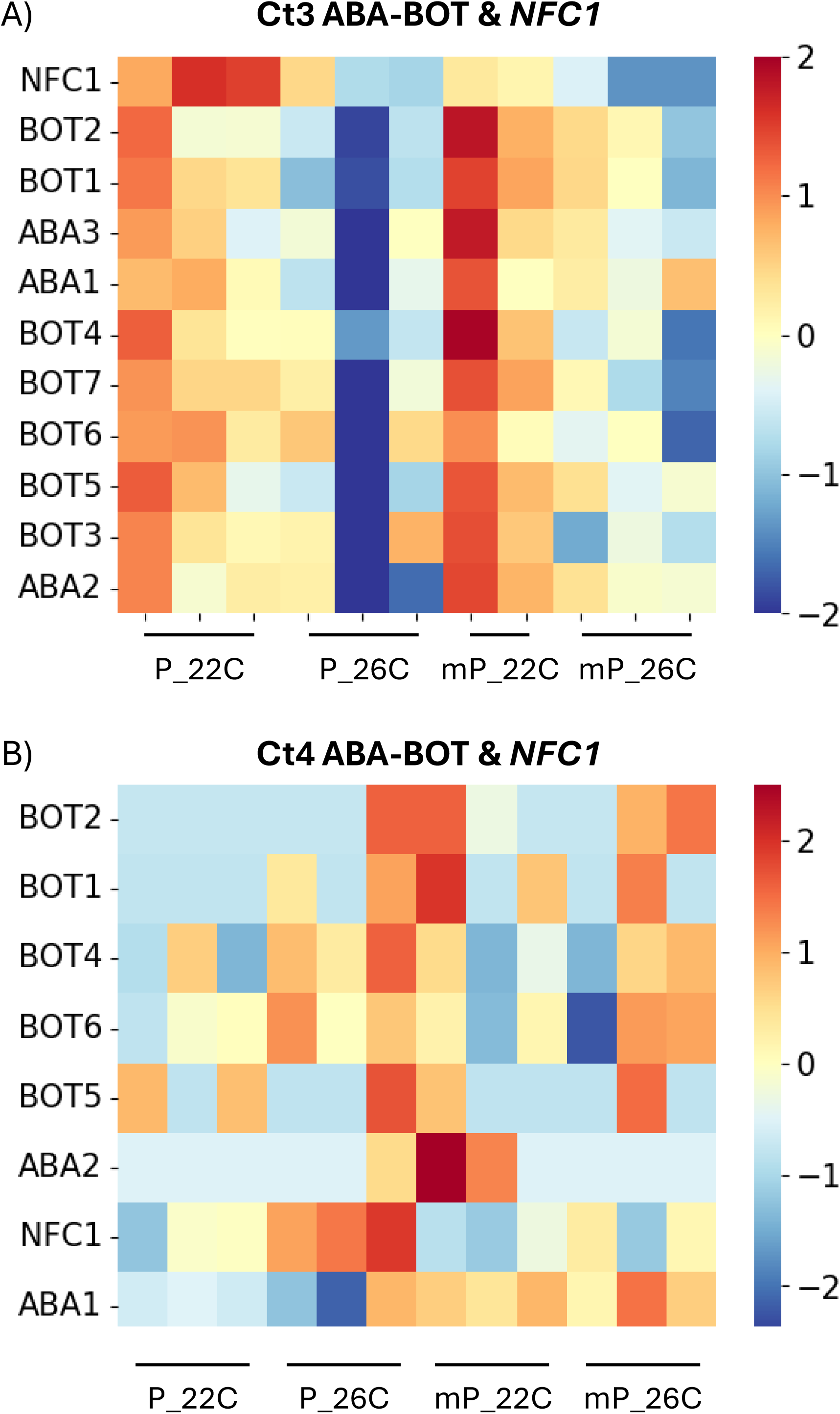
Ct3 and Ct4 have contrasting virulence factor expression profiles. A, B) Heatmap of ABA-BOT genes and *NFC1* in A) Ct3, B) Ct4 inoculated to Arabidopsis at 26°C or 22°C under Pi sufficient (P) or deficient conditions (mP).

**Figure S3.**
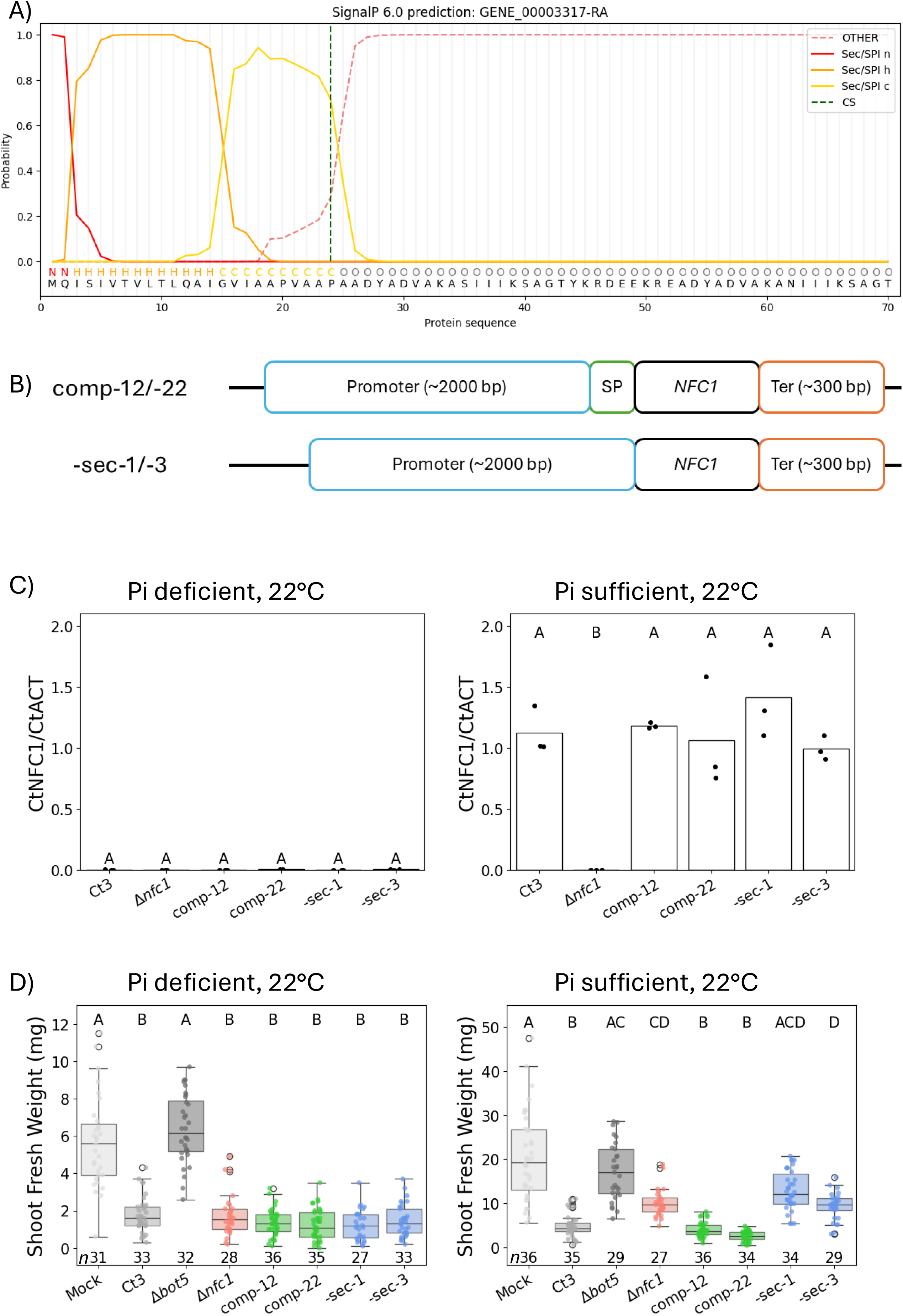
*NFC1* has a secretion signal necessary for its function in virulence. A) SignalP prediction of the signal peptide in *NFC1* (GENE_00003317-RA). Green dashed line represents the predicted cleavage site (CS). B) Diagram depicting full complementation lines (comp-12/-22) and complementation lines without the predicted signal peptide (-sec-1/-3). SP (outlined in green) represents the predicted signal peptide, and Ter (outlined in orange) represents the terminator. C) Relative expression of *NFC1* in fungi inoculated to plants at 22°C at 2 dpi after direct inoculation. Letters above each bar represent significance levels from a post-hoc Tukey’s HSD test (*p* < 0.05). D) Shoot fresh weight of plants inoculated with fungi at 22°C at 24 dpi from seed inoculation. Letters above each box represent significance levels from a Dunn test (*p* < 0.05). Numbers below each box represent sample size.

**Figure S4.**
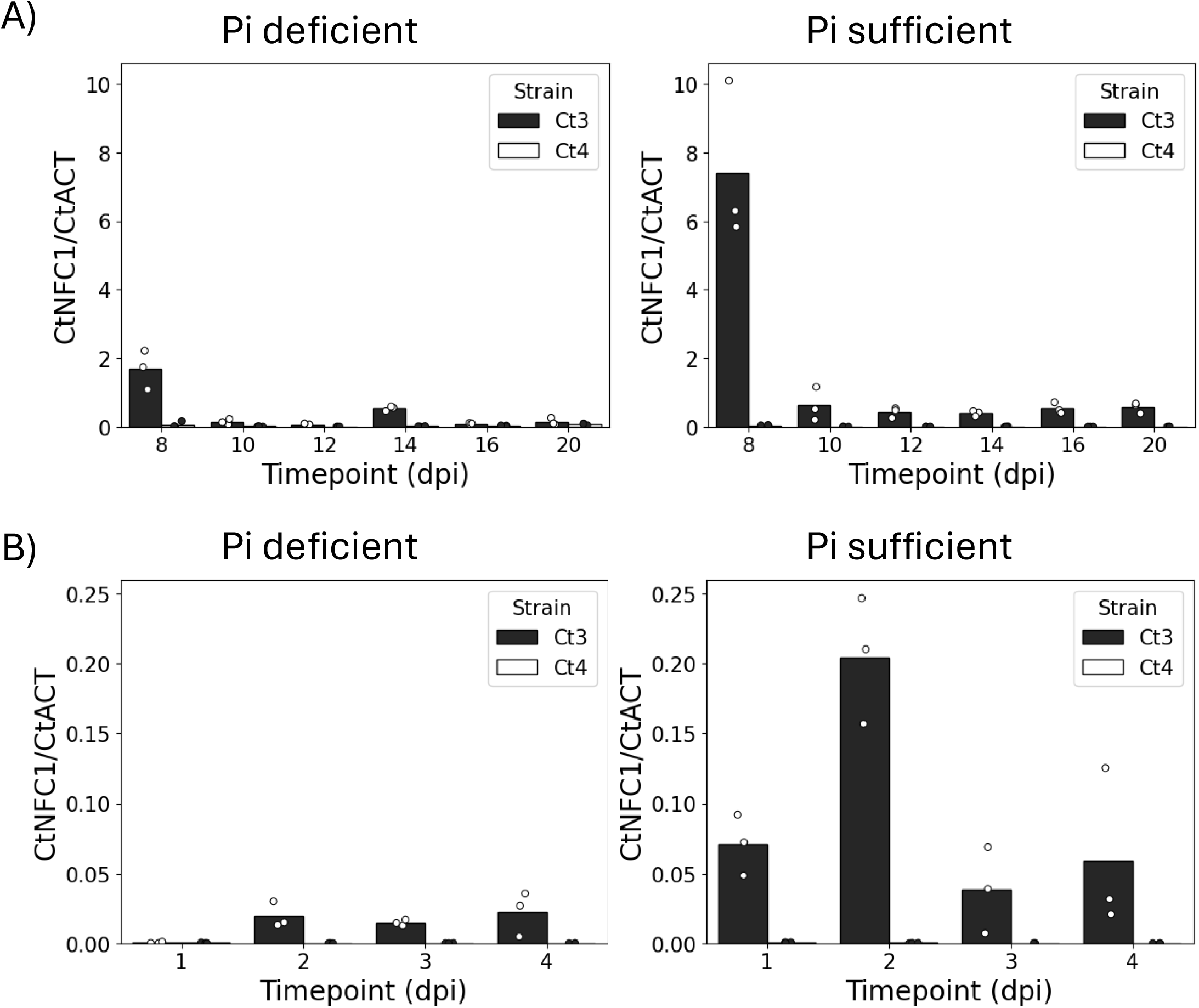
*NFC1* is expressed in the early timepoint under Pi sufficient conditions at 22°C. A, B) Relative expression of *NFC1* normalized to *CtACTIN* from A) 8 to 20 dpi after seed inoculation, B) 1 to 4 dpi after direct inoculation at 22°C under Pi sufficient or deficient conditions.

**Figure S5.**
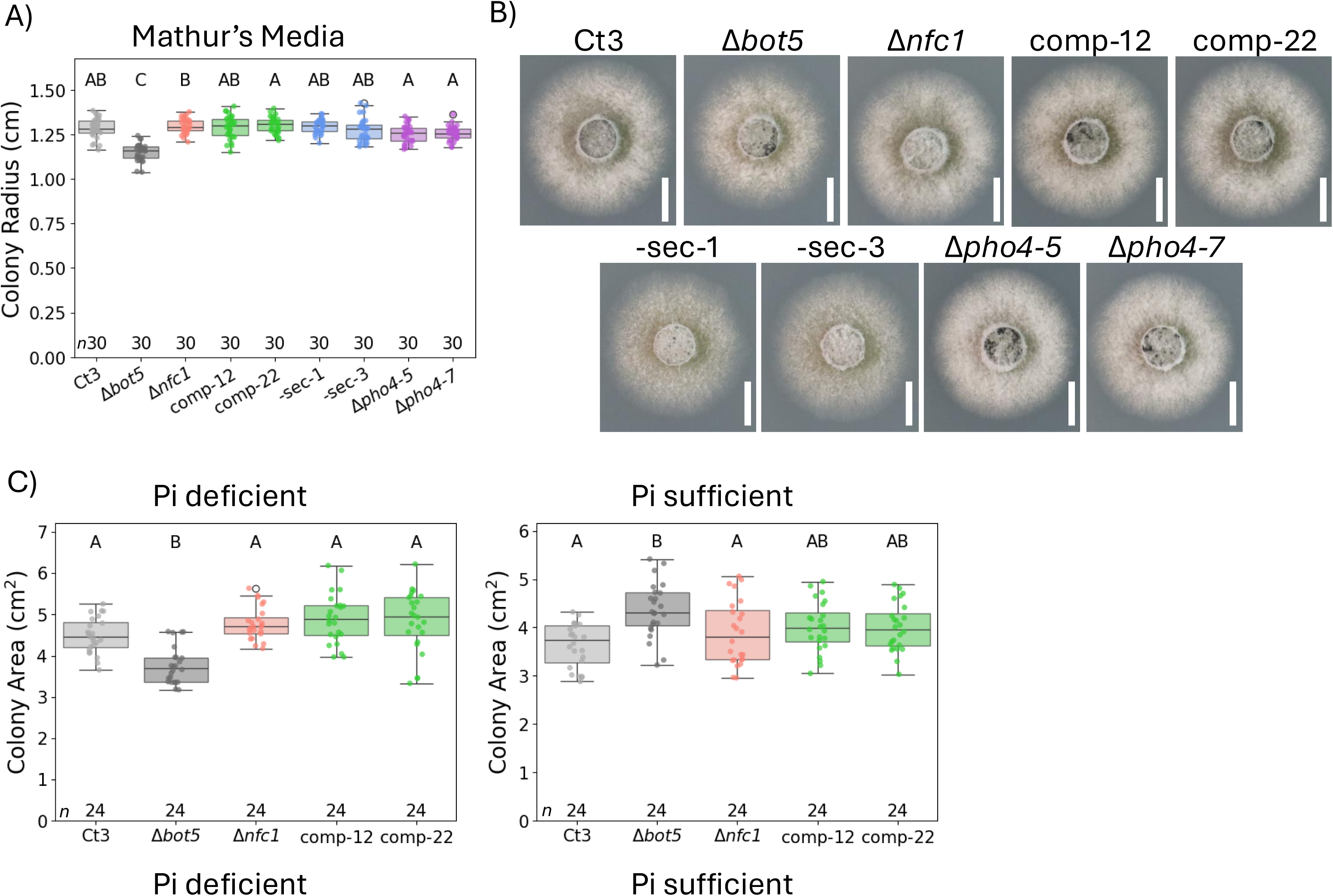
Neither Δ*nfc1* nor Δ*pho4* show growth deficits on rich media. A) Colony radius of colonies grown from 5 mm agar plugs for 3 days on Mathur’s media plates. Letters above each box represent significance levels from a Dunn test (*p* < 0.05). Numbers below each box represent sample size. B) Representative images of colonies quantified in (A). Scale bars represent 1 cm. C) Colony area of colonies grown from spores for 10 days on modified half-strength MS plates with deficient or sufficient Pi at 22°C. Letters above each box represent significance levels from a Dunn test (*p* < 0.05). Numbers below each box represent sample size.

**Figure S6.**
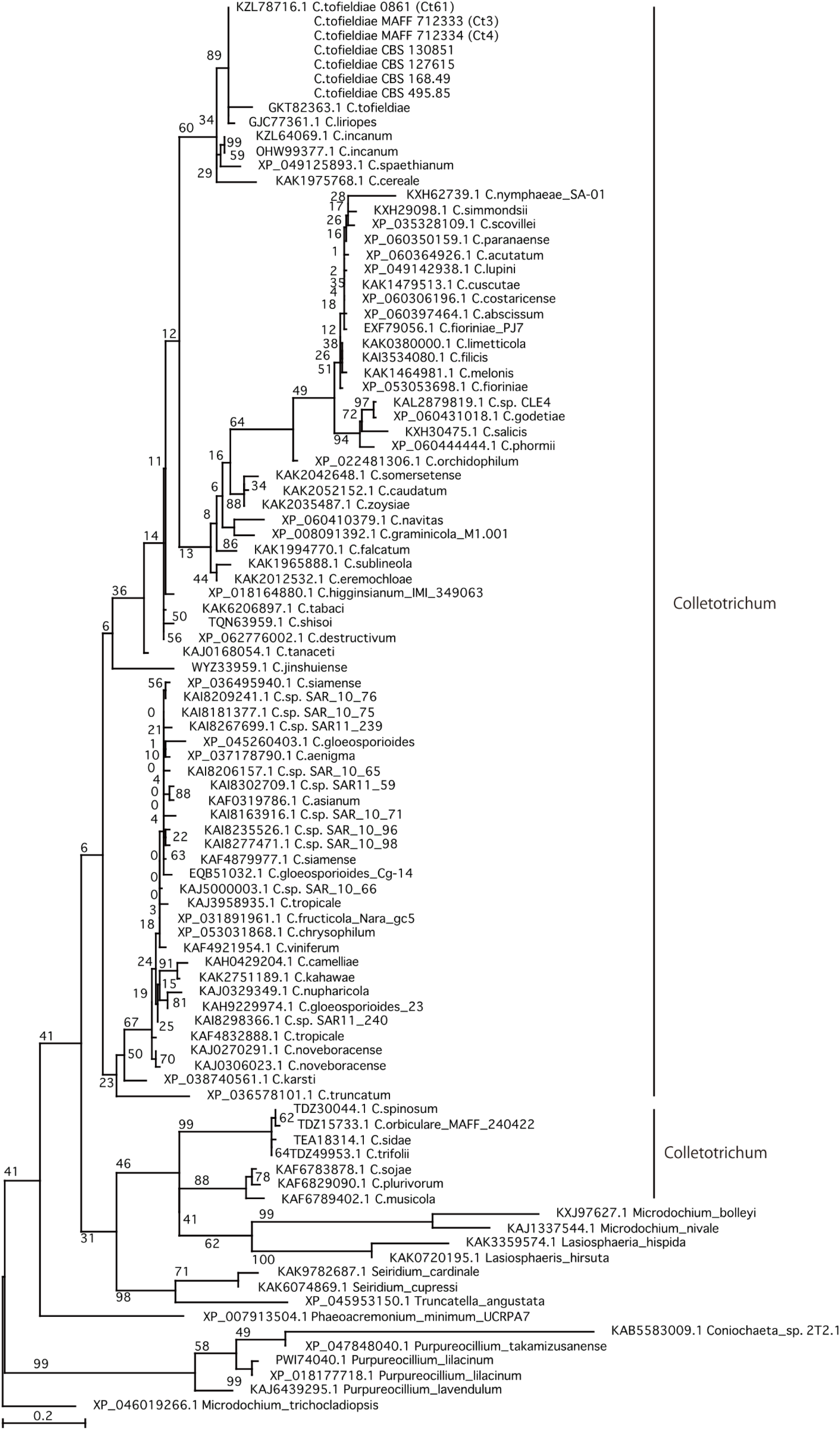
*NFC1* is a phylogenetically conserved gene. Maximum likelihood phylogenetic tree based on the amino acid sequence of NFC1 retrieved from NCBI WGS contigs and the nr protein database.

**Figure S7.**
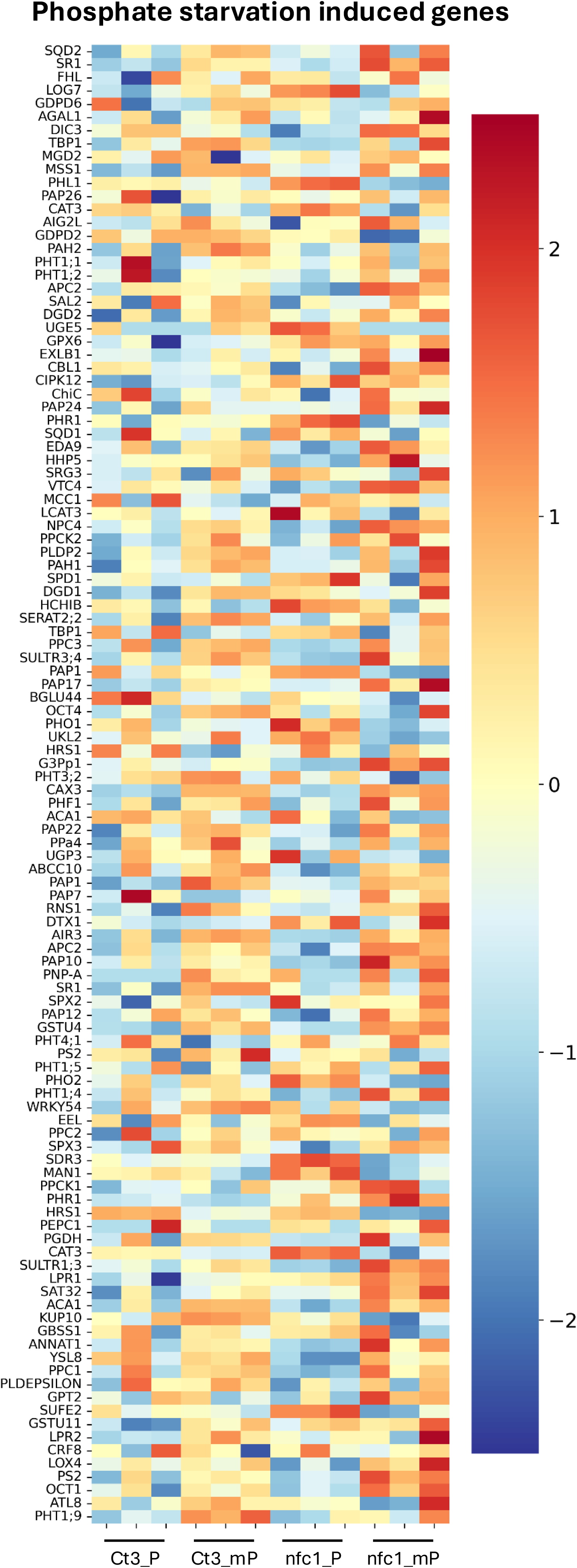
*NFC1* does not induce PSI genes at 2 dpi. Heatmap of Arabidopsis PSI genes in response to Ct3 or Δ*nfc1* inoculation under Pi sufficient (P) or Pi deficient (mP) conditions.

**Figure S8.**
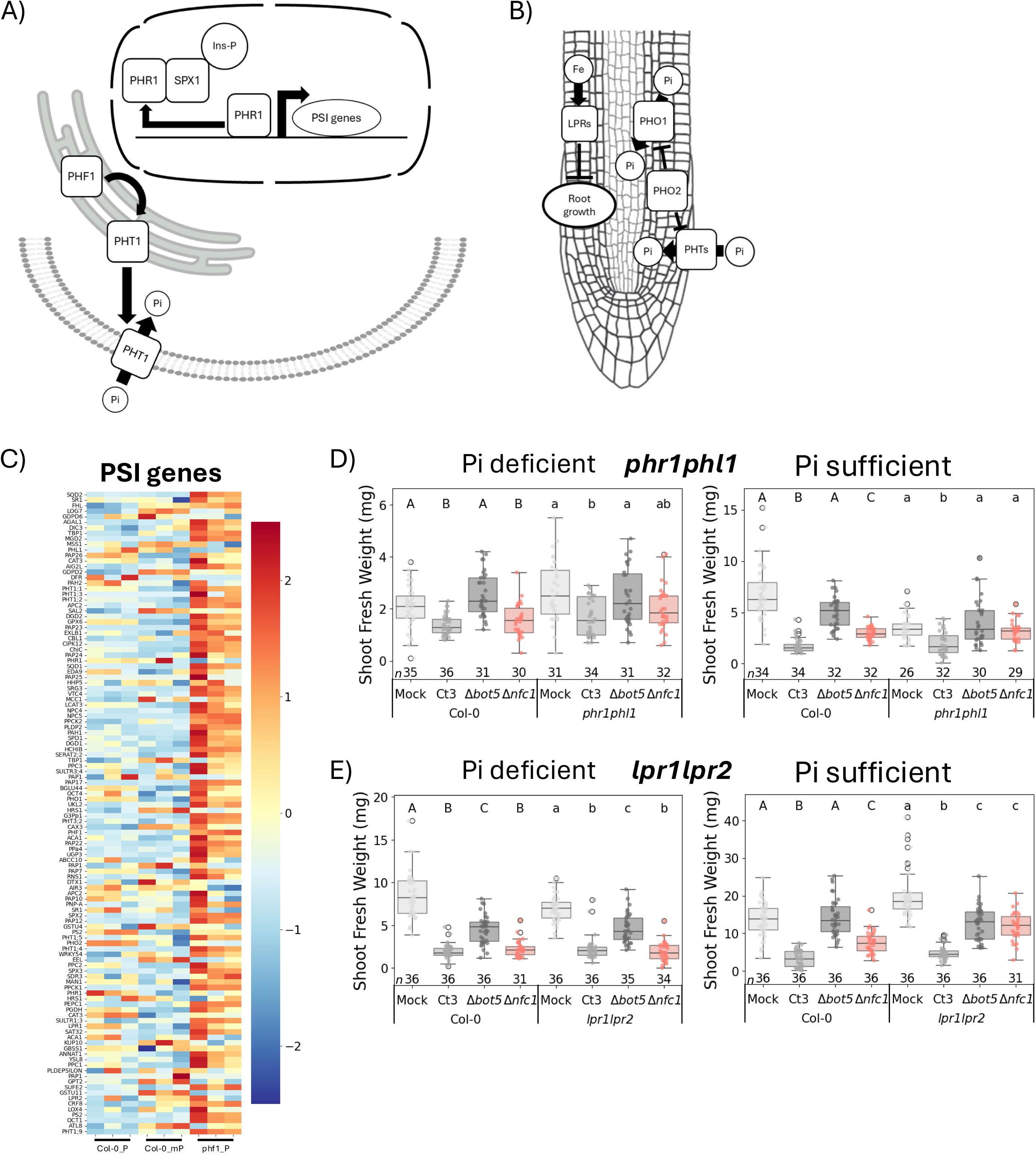
*NFC1* does not depend on root phosphate sensing or the PSR. A, B) Schematic diagrams of key PSR components at a A) cellular level and B) root organ level. A) Under Pi deficient conditions, PHF1 allows phosphate transporters (e.g. PHT1 family) to be trafficked from the endoplasmic reticulum to the plasma membrane to transport Pi into the cell. At the same time, SPX1-domain-containing proteins do not bind with Ins-PPs and therefore do not bind PHR1, allowing it to bind to DNA and activate expression of phosphate starvation-induced (PSI) genes. B) Under Pi deficient conditions, there is a relatively higher amount of iron, activating LPR1/LPR2, which inhibits primary root growth. At the same time, PHO2 is inhibited to release degradation of phosphate transporters (e.g. PHTs that transport Pi into the root or PHO1 that transports Pi into the xylem). C) Heatmap of Mock-inoculated plants at 2 dpi demonstrating that PSI genes are induced in *phf1* plants. D, E) Shoot fresh weight of D) *phr1phl1*, E) *lpr1/2* plants inoculated with fungi at 22°C at 17 dpi from seedling inoculation. Letters above each box represent significance levels from a Dunn test (*p* < 0.05). Numbers below each box represent sample size.

**Figure S9.**
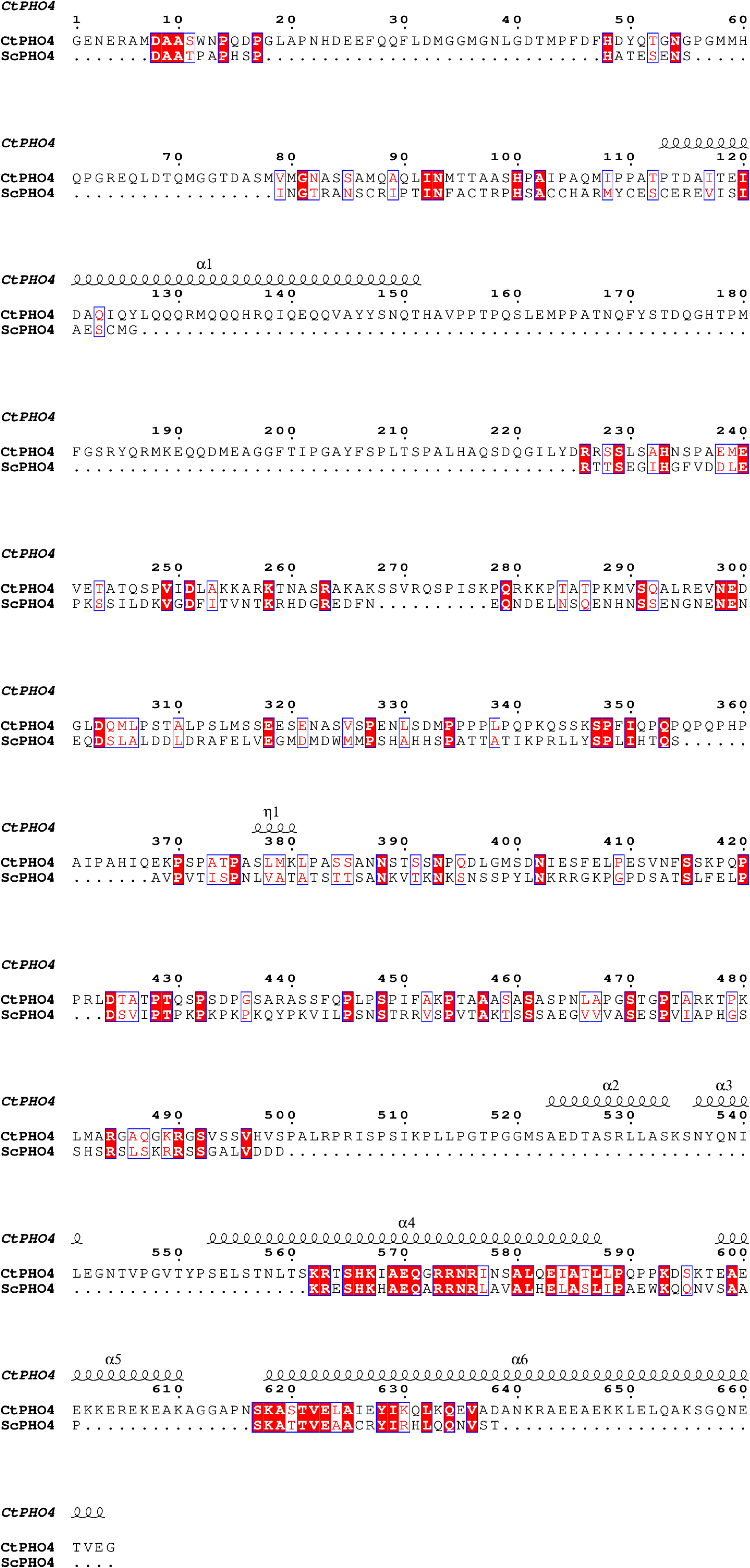
CtPHO4 is an orthologue of ScPHO4. Sequence alignment of amino acid sequences of CtPHO4 (top) and ScPHO4 (bottom). Highlighted in red (white text) and outlined in blue are identical amino acids in CtPHO4 and ScPHO4, and highlighted in white (red text) and outlined in blue are group similarity amino acids. Structural elements of CtPHO4 based on AlphaFold3 structural prediction are represented above the sequences.

**Figure S10.**
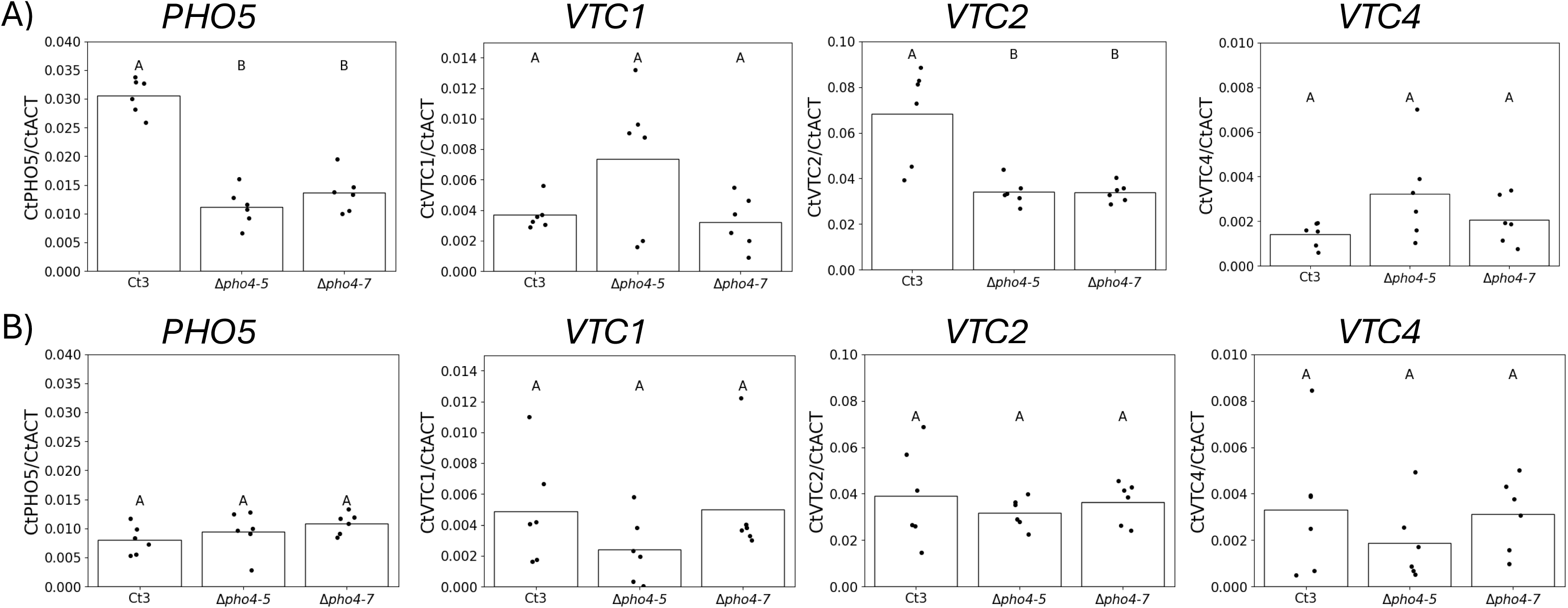
*CtPHO4* is a PSR regulator in Ct3. A, B) Relative expression of *CtPHO5* and *CtVTC*s under A) Pi deficient, B) Pi sufficient conditions from mycelia grown in liquid half-strength MS with sucrose for 5 days at room temperature. Letters above each bar represent significance levels from a post-hoc Tukey’s HSD test.

## Notes

### Competing Interest Statement

The authors have declared no competing interest.

